# Glucose intolerance in aging is mediated by the Gpcpd1-GPC metabolic axis

**DOI:** 10.1101/2021.10.26.465828

**Authors:** Domagoj Cikes, Michael Leutner, Shane J.F. Cronin, Maria Novatchkova, Lorenz Pfleger, Radka Klepochová, Benjamin Lair, Marlène Lac, Camille Bergoglio, Nathalie Viguerie, Gerhard Dürnberger, Elisabeth Roitinger, Eric Rullman, Thomas Gustafsson, Astrid Hagelkruys, Geneviève Tavernier, Virginie Bourlier, Claude Knauf, Michael Krebs, Alexandra Kautzky-Willer, Cedric Moro, Martin Krssak, Michael Orthofer, Josef M. Penninger

## Abstract

Skeletal muscle plays a central role in the regulation of systemic metabolism during lifespan. With aging, muscle mediated metabolic homeostasis is perturbed, contributing to the onset of multiple chronic diseases. Our knowledge on the mechanisms responsible for this age-related perturbation is limited, as it is difficult to distinguish between correlation and causality of molecular changes in muscle aging. Glycerophosphocholine phosphodiesterase 1 (GPCPD1) is a highly abundant muscle enzyme responsible for the hydrolysis of the lipid glycerophosphocholine (GPC). The physiological function of GPCPD1 remained largely unknown. Here, we report that the GPCPD1-GPC metabolic pathway is dramatically perturbed in the aged muscle. Muscle-specific inactivation of *Gpcpd1* resulted in severely affected glucose metabolism, without affecting muscle development. This pathology was muscle specific and did not occur in white fat-, brown fat- and liver-deficient *Gpcpd1* deficient mice. Moreover, in the muscle specific mutant mice, glucose intolerance was markedly accelerated under high sugar and high fat diet. Mechanistically, *Gpcpd1* deficiency results in accumulation of GPC, without any other significant changes in the global lipidome. This causes an “aged-like” transcriptomic signature in young *Gpcpd1* deficient muscles and impaired insulin signaling. Finally, we report that GPC levels are markedly perturbed in muscles from both aged humans and patients with Type 2 diabetes, with a high correlation between GPC levels and increased chronological age. Our findings show the novel and critical physiological function of GPCPD1-GPC metabolic pathway to glucose metabolism, and the perturbation of this pathway with aging, which may contribute to glucose intolerance in aging.

## Introduction

Skeletal muscle is the biggest organ in the body, essential for mobility, health-span and lifespan (Cruz-Jentoft and Sayer, 2019; Yanagi et al., 2021; Yang et al., 2019). Apart from mobility, muscle plays a crucial role in systemic metabolism (Baskin et al., 2015). Critical to this regulatory role is the control of glucose metabolism, with muscle metabolizing up to 80% of ingested glucose (DeFronzo et al., 1981; Jue et al., 1989). Aged muscles exhibit multiple molecular and metabolic perturbations, that can potentially affect muscle function and whole-body metabolism during the lifespan (Demontis et al., 2013a). With muscle ageing, glucose utilization is significantly impaired, leading to systemic perturbations of glucose metabolism (Chia et al., 2018; Kern et al., 1992). This systemic metabolic dysfunction contributes to the onset and progression of multiple chronic diseases (Cowie et al., 2009).

Glycerophosphodiester phosphodiesterase 1 (Gpcpd1; Gdpd6; Edi3) is a member of a large family of glycerophosphodiester phosphodiesterases, highly conserved enzymes found in bacteria, protozoa, and mammals (Corda et al., 2014). Mammalian glycerophosphodiesterases exist in multiple isoforms (seven in humans) with a high degree of tissue and substrate specificity (Corda et al., 2014). Gpcpd1 is a 76.6 kD protein that hydrolyzes lipid glycerophosphocholine (GPC), yielding choline and glycerol-3-phosphate. Unlike other phosphodiesterases, Gpcpd1 does not contain a transmembrane region and is localized in the cytoplasm. An early study reported that Gpcpd1 expression is enriched in heart and skeletal muscle (Okazaki et al., 2010). The same study also reported that over-expression of a truncated version of Gpcpd1 in the muscles resulted in muscle atrophy, suggesting that Gpcpd1 regulates muscle differentiation (Okazaki et al., 2010). High expression of Gpcpd1 was found in metastatic endometrial cancers, while inhibition of Gpcpd1 in breast cancer cells inhibited migration and invasion (Stewart et al., 2012). Apart from these studies, the physiological function of Gpcpd1 remains largely unknown, and an *in vivo* loss-of-function analysis has never been reported.

A recent untargeted metabolomic profiling of skeletal muscle from old mice indicated glycerophosphocholine (GPC), the Gpcpd1 substrate, as a significantly elevated metabolite in aged mouse muscles (Houtkooper et al., 2011). Moreover, a single nucleotide polymorphism in proximity to the human GPCPD1 locus was found to be associated with longevity (Pilling et al., 2016). Therefore, we set out to investigate if there is a direct link between the Gpcpd1-GPC metabolic pathway and aging-related health decline.

## MATERIALS AND METHODS

### Animal care

All experimental protocols were approved by the institutional Animal Ethics Committee in accordance with the Austrian legal guidelines on Animal Care. All mice were on a C57BL/6J background (in-house colony). *MckCre, Ap2Cre, Ucp1Cre, AlbCre* mice were purchased from the Jackson Laboratory (Bar Harbor, US, stock number 006405, 005069, 024670, 003574). The mice were maintained on a 12:12 hour light-dark cycle (lights on 08:00-20:00) while housed in ventilated cages at an ambient temperature of 25 °C. Mice were fed *ad libitum* standard chow diet (17% kcal from fat; Envigo, GmbH), high-fat diet (HFD; 60% kcal from fat; Envigo, GmbH), or high fructose diet (30% fructose in drinking water; #F0127 Sigma Aldrich Gmbh) starting from 8 weeks of age.

### Human biopsies

All human experiments were approved by the regional ethical review board in Stockholm (2014/516-31/2 and 2010/786-31/3) and complied with the Declaration of Helsinki. Oral and written informed consent were obtained from all subjects prior to participation in the study. 8 healthy young adults (age 21-29) and 8 middle-aged (age 45-62) subjects were recruited. The subjects did not use any medications and were nonsmokers. Biopsies of the quadriceps skeletal muscle (vastus lateralis) were obtained under local anesthesia using the Bergström percutaneous needle biopsy technique (Bergström and Hultman, 1967). The biopsies were immediately frozen in isopentane, cooled in liquid nitrogen, and stored at −80°C until further analysis.

### Determination of muscle glycerophosphocholine in Type 2 diabetes patients

This prospective clinical study was performed at the Department of Internal Medicine 3, Division of Endocrinology and Metabolism at the Medical University of Vienna between June 2021 and September 2021. Patients were recruited at the Diabetes Outpatient Department of the Medical University of Vienna and enrolled for the study after giving written informed consent. Routine laboratory measurements were analyzed at the certified Department of Medical and Chemical Laboratory Diagnostics (http://www.kimcl.at/) of the Medical University of Vienna. In the present study, we included patients with diagnosed diabetes mellitus type 2 and patients without diabetes mellitus aged between 50 and 65 years. Exclusion criteria were infectious diseases such as hepatitis B or C or human immunodeficiency virus (HIV) We performed a detailed metabolic characterization, including measurements of lipid parameters, glucose metabolism (e.g. HbA1c, fasting glucose levels), routine laboratory analyses and anthropometric measurements. *In vivo* magnetic resonance spectroscopy (MRS) of Vastus lateralis muscle was performed on individuals that were selected based on their age and without the use of any medications or a record of musculoskeletal or cardiovascular disease (Krumpolec et al., 2020; Valkovič et al., 2013). Laboratory analyses and MRS measurements were performed after a 12-hour fasting period. The study was approved by the ethics board of the Medical University of Vienna.

### Cross tissue Gpcpd1 expression analysis

The data on Gpcpd1 mRNA expression analysis in several tissues were derived from the gene expression profiling atlas GSE10246 (Lattin et al., 2008) and visualized using the BioGPS portal (Wu et al., 2016).

### Generation of *Gpcpd1* conditional and tissue specific mutant mice

*Gpcpd1* conditional knockout mice were derived from targeted ES cells obtained from EUCOMM (European Conditional Mouse Mutagenesis Program). Exons 9 and exon 10 of the *Gpcpd1* gene were flanked by loxP sites. Cre mediated deletion of the floxed region creates a frameshift and truncated protein. Upon confirmation of correct targeting by Southern blotting, the ES cell clone G8 (C57Bl/6N) was injected into C57BL/6J-Tyr ^c-2J^/J blastocysts and offspring chimeric mice were crossed to C57BL/6J mice. Following germline transmission, targeted mice were crossed to transgenic mice expressing FLPe recombinase leading to excision of the NEO cassette. Offspring mice were backcrossed onto a C57Bl/6J genetic background. Following primers were used to identify the floxed allele:

Forward – GTGCAGGGAACTCAACAACG

Reverse - AGTGATGACAAAGAGGCCAAAAAG

Subsequently, *MckCre, Ap2Cre, Ucp1Cre,* and *AlbCre* mutant animals were crossed to the *Gpcpd1*^flox/flox^ mice to generate *MckCre-Gpcpd1, Ap2Cre-Gpcpd1, Ucp1Cre-Gpcpd1,* and *AlbCre-Gpcpd1* mutant mice.

### Mass spectrometry

25μl of each sample containing 100μg protein in RIPA buffer were mixed with 25 μl of 2x lysis buffer (iST, PO 00065, PreOmics) and sample preparation and tryptic digest were performed using the iST 96x kit (PO 00027, PreOmics) according to the manufacturer’s description.

750ng of each generated peptide sample was analysed by nanoLC-MS/MS. The nano HPLC system (UltiMate 3000 RSLC nano system, Thermo Fisher Scientific) was coupled to an Exploris 480 mass spectrometer equipped with a FAIMS pro interfaces and a Nanospray Flex ion source (all parts Thermo Fisher Scientific). Peptides were loaded onto a trap column (PepMap Acclaim C18, 5 mm × 300 μm ID, 5 μm particles, 100 Å pore size, Thermo Fisher Scientific) at a flow rate of 25 μl/min using 0.1% TFA as mobile phase. After 10 minutes, the trap column was switched in line with the analytical column (a prototype 110 cm μPAC™ Neo HPLC column, Thermo Fisher Scientific) operated at 30°C. Peptides were eluted using a flow rate of 300 nl/min, starting with the mobile phases 98% A (0.1% formic acid in water) and 2% B (80% acetonitrile, 0.1% formic acid) and linearly increasing to 35% B over the next 180 minutes.

The Exploris mass spectrometer was operated in data-dependent mode, performing a full scan (m/z range 350-1200, resolution 60,000, target value 1E6) at 3 different compensation voltages (CV -45, -60, -75), each followed by MS/MS scans of the most abundant ions for a cycle time of 0.8 sec per CV. MS/MS spectra were acquired using a collision energy of 30, isolation width of 1.0 m/z, resolution of 30.000, target value of 1E5 and intensity threshold of 2.5E4, maximum injection time was set to 30 ms. Precursor ions selected for fragmentation (include charge state 2-6) were excluded for 40 s. The monoisotopic precursor selection filter and exclude isotopes feature were enabled. For peptide identification, the RAW-files were loaded into Proteome Discoverer (version 2.5.0.400, Thermo Scientific). All MS/MS spectra were searched using MSAmanda v2.0.0.16129 (Dorfer V. et al., J. Proteome Res. 2014 Aug 1;13(8):3679-84). The peptide and fragment mass tolerance was set to ±10 ppm, the maximum number of missed cleavages was set to 2, using tryptic enzymatic specificity without proline restriction. Peptide and protein identification was performed in two steps. For an initial search the RAW-files were searched against the database uniprot_reference_mouse_2022-03-04.fasta (21,962 sequences; 11,728,099 residues), supplemented with common contaminants and sequences of tagged proteins of interest, using the following search parameters: Iodoacetamide derivative on cysteine was set as a fixed modification, oxidation of methionine as variable modification. The result was filtered to 1 % FDR on protein using the Percolator algorithm (Käll L. et al., Nat. Methods. 2007 Nov; 4(11):923-5) integrated in Proteome Discoverer. A sub-database of proteins identified in this search was generated for further processing. For the second search, the RAW-files were searched against the created sub-database using the same settings as above plus considering additional variable modifications: Phosphorylation on serine, threonine and tyrosine, deamidation on asparagine and glutamine, and glutamine to pyro-glutamate conversion at peptide N-terminal glutamine, acetylation on protein N-terminus were set as variable modifications. The localization of the post-translational modification sites within the peptides was performed with the tool ptmRS, based on the tool phosphoRS (Taus T. et al., J. Proteome Res. 2011, 10, 5354-62). Identifications were filtered again to 1 % FDR on protein and PSM level, additionally an Amanda score cut-off of at least 150 was applied. Peptides were subjected to label-free quantification using IMP-apQuant (Doblmann J. et al., J. Proteome Res. 2019, 18(1):535-541). Proteins were quantified by summing unique and razor peptides and applying intensity-based absolute quantification (iBAQ, Schwanhäusser B. et al., Nature 2011, 473(7347):337−42) with subsequent normalisation based on the MaxLFQ algorithm (Cox J. et al., Mol Cell Proteomics. 2014, 13(9):2513-26). Identified proteins were filtered to contain at least 3 quantified peptide groups. Protein-abundances were normalized using sum normalization. Ion chromatograms extracted by apQuant were plotted using R.

### Quantitative real-time PCR (RT-qPCR)

After animal sacrifice, tissues were extracted, separated into regions and flash frozen in liquid nitrogen. Subsequently, mRNA was isolated using the RNeasy Lipid Tissue Mini Kit (Qiagen, GmbH). The RNA concentrations were estimated by measuring the absorbance at 260 nm using Nanodrop (Thermofisher, GmbH). cDNA synthesis was performed using the iScript Advanced cDNA Synthesis Kit for RT-qPCR (Bio-Rad, GmbH) following manufacturer’s recommendations. cDNA was diluted in DNase-free water (1:10) before quantification by real-time PCR. mRNA transcript levels were measured in triplicate samples per animal using CFX96 touch real-time PCR (Bio-Rad GmbH). Detection of the PCR products was achieved with SYBR Green (Bio-Rad, GmbH). At the end of each run, melting curve analyses were performed, and representative samples of each experimental group were run on agarose gels to ensure the specificity of amplification. Gene expression was normalized to the expression level of 18S ribosomal rRNA as the reference gene. The following primers were used:

Murine Gpcpd1:

Forward 5’- GTGGTGCAGGGAACTCAACAACG - 3’

Reverse 5’- TGAGGTCATGATACACCACGGGC - 3’

Murine 18SrRNA:

Forward 5’- GGCCGTTCTTAGTTGGTGGAGCG -3’

Reverse 5’- CTGAACGCCACTTGTCCCTC - 3’

Human Gpcpd1:

Forward 5’- GCATCTGTGGTGCTAGGTGA - 3’

Reverse 5’- TGCCTTGTGAAAAACATGCAG - 3’

Human 18SrRNA:

Forward 5’- GGCCCTGTAATTGGAATGAGTC -3’

Reverse 5’- CCAAGATCCAACTACGAGCTT - 3’

### Grip strength analysis

20 month old mice (control and Pcyt2 Myf5 KO mice) were subjected to grip strength tests using a grip strength meter (Bioseb, USA), following standardized operating procedures. Prior to tests, mice were single caged for two weeks, in order to avoid any littermate influence on their performance. Each mouse was tested three times, with a 15-minute inter-trial interval, and values were averaged among the three trials. Clasping index was evaluated as described previously 84. Each mouse was scored three times, and an average of scores was calculated.

### Glucose tolerance test (GTT), insulin tolerance test (ITT) and insulin measurements

For glucose tolerance tests (GTT), mice were fasted for 16 h and a D-glucose solution was administered by oral gavage at a dose of 2 g/kg (chow diet), or 1 g/kg (high sugar, high fat diet). Blood samples were collected from the tail vein and glucose was measured using a glucometer (Roche, Accu-Chek Performa). For analysis of insulin levels, tail vein blood samples were added to a NaCl/EDTA solution to avoid blood clotting and plasma insulin concentrations were determined using the Alpco Mouse Ultrasensitive Insulin ELISA (80-INSMSU-E10). For insulin tolerance tests (ITT), mice were fasted for 6 h and insulin solution (0.375 IU insulin/kg) administered i.p., followed by blood glucose measurements.

### Radioactive 2-deoxyglucose-6-phosphate tissue uptake measurements

To determine 2-DG uptake in various tissues, a bolus injection of 2-[1,2-3H(N)]deoxy-D-glucose (PerkinElmer, Boston, Massachusetts) (0.4 μCi/g body weight) and insulin (3mU/g body weight) was injected intraperitoneally in mice (Laurens et al., 2016a, 2016b). Mice were killed 30 min after injection and tissues were snap-frozen and stored at -80°C until further processing. Tissue-specific accumulation of 2-deoxyglucose-6-phosphate was determined as previously described with minor modifications (Kraegen et al., 1985). The total of the [3H]-radioactivity found in 2-deoxyglucose-6-phosphate was divided by the mean specific activity of glucose at 30 minutes to obtain the tissue-specific metabolic clearance index (Rg) (μmol per 100 g of wet tissue per minute).

### C2C12 myotubes glucose uptake

C2C12 were seeded in 24-well dishes (Corning, Sigma-Aldrich, GmbH) and grown in DMEM/F12 with 10% FCS and penicillin–streptomycin (Life Technologies, final conc., 50 U/mL of penicillin). Differentiation was achieved by culturing the cells in differentiation media (DMEM/F12 with 2% horse serum and penicillin–streptomycin), after achieving 80% confluency. 48 hours after differentiation, in the experiment group differentiation media was supplemented with 10mM glycerophosphocholine (#20736, Cayman Chemicals, USA). Cells were incubated further in differentiation media with or without glycerophosphocholine for additional 7 days, with daily medium changes and intracellular glycerophosphocholine levels were determined using mass spectrometry as stated above. On the day of the glucose uptake measurements, differentiated myotubes were washed twice with Hank’s Balanced Salt Solution (Gibco) (HBSS) and further starved in HBSS with or without glycerophosphocholine (10mM) for 6h. Afterwards, cells were washed twice with HBSS, and incubated with HBSS containing only D-glucose (15mM) and 200nM insulin (#I0516, Sigma Aldrich, GmbH) for 20 minutes. The solution was washed after indicated time with HBSS, the reaction was stopped by adding 0.6N HCl, neutralized with 1M Tris solution and lysates were further processed for measurement using the Glucose-Glo Assay (Promega, GmbH) according to the manufacturer’s instructions.

### Determination of muscle glycogen levels

Skeletal muscle tissue (quadriceps) was surgically removed, and flash frozen in liquid nitrogen. The tissue was crushed on dry ice, weighed, and further processed according to the manufacturers’ instructions (Sigma Aldrich; #MAK016). Tissue glycogen levels were determined based on the formulation derived from the standard curve and normalized to the tissue weight.

### Tissue RNA purification

Total RNA was extracted from skeletal muscles using the Qiagen miRNeasy Mini kit (cat# 217004, Qiagen, GmbH) per the manufacturer’s instructions. In brief, samples were homogenized in QIAzol lysis reagent using a rotor stator homogenizer, and then centrifuged after chloroform addition at 12,000⍰g to separate the organic and aqueous phases. Total RNA was purified from the aqueous phase using the spin column provided in the kit. DNA was digested on-column per the manufacturer’s instructions. RNA concentration was measured using the Nanodrop and RNA integrity was measured with an Agilent 2200 Tapestation instrument (cat#5067–5576 and cat#5067–5577, Agilent, Santa Clara, CA). All samples had RIN values greater than 8.

### Quantseq analysis and sequencing

Libraries were prepared using the QuantSeq 3’mRNA-Seq Library Prep Kit-FWD (cat #15, Lexogen, Vienna, Austria) using 1⍰μg of RNA per library, following manufacturer’s instructions.

11⍰cycles of library amplification were performed with indices from the first two columns of the i7 Index Plate for QuantSeq/SENSE with Illumina adapters 7001–7096 (cat #044, Lexogen). Libraries were eluted in 22⍰μL of the Elution Buffer, and double stranded DNA concentration was quantified by the KAPA Library Quantification Kit. The molar concentration of cDNA molecules was calculated from the double stranded DNA concentration and the region average size (determined by analyzing each sample on an Agilent 2200 Tapestation instrument (cat#5067–5584 and cat#5067–5585, Agilent, Santa Clara, CA)). Aliquots containing an equal number of nmoles of cDNA molecules from each library were pooled to a concentration of 10⍰nM of cDNA. The final pool was purified once more (to remove any free primers to prevent index-hopping) by adding 0.9x volumes of PB and proceeding from Step 30 onwards in the QuantSeq User Guide protocol. The library was eluted in 22⍰μL of the kit’s Elution Buffer. The pooled libraries were sequenced using an Illumina HiSeq4000 instrument (Illumina, San Diego, CA).

### Transcript coverage

RNA-seq reads were trimmed using BBDuk v38.06 (ref=polyA.fa.gz,truseq.fa.gz k=13 ktrim=r useshortkmers=t mink=5 qtrim=r trimq=10 minlength=20) and reads mapping to abundant sequences included in the iGenomes Ensembl GRCm38 bundle (mouse rDNA, mouse mitochondrial chromosome, phiX174 genome, adapter) were removed using bowtie2 v2.3.4.1 alignment. Remaining reads were analyzed using genome and gene annotation for the GRCm38/mm10 assembly obtained from Mus musculus Ensembl release 94. Reads were aligned to the genome using star v2.6.0c and reads in genes were counted with featureCounts (subread v1.6.2) using strand-specific read counting for QuantSeq experiments (-s 1). Differential gene expression analysis on raw counts and variance-stabilized transformation of count data for heatmap visualization were performed using DESeq2 v1.18.1. Functional annotation overrepresentation analysis of differentially expressed genes was conducted using clusterprofiler v3.6.0 in R v3.4.1.

### Western blot analysis and immunofluorescence

A D-glucose solution was administered by oral gavage and mice were sacrificed after 15 minutes. Muscle tissue (quadriceps) was immediately surgically removed, and flash frozen in liquid nitrogen. Tissues were further homogenized in RIPA buffer (Sigma; R0278) containing Halt protease and phosphatase inhibitor cocktail (Thermo Scientific; 78440). Protein levels were determined using the Bradford assay kit (Pierce, GmbH) and lysates containing equal amounts of protein were subjected to SDS-PAGE, further transferred to nitrocellulose membranes. Western blotting was carried out using standard protocols. Blots were blocked for 1 hour with 5% BSA (Sigma Aldrich; #820204) in TBST (1× TBS and 0.1% Tween-20) and were then incubated overnight at 4°C with primary antibodies diluted in 5% BSA in TBST (1:250 dilution). Blots were washed three times in TBST for 15 min, then incubated with HRP-conjugated secondary antibodies diluted in 5% BSA in TBST (1:5000 dilution) for 45 min at room temperature, washed three times in TBST for 15 min and visualized using enhanced chemiluminescence (ECL Plus, Pierce, 1896327). The following primary antibodies were used: anti-phospho-Insulin Receptor β (Tyr1150/1151) (# 3021 CST, DE, 1:250), anti-total Insulin Receptor β (# 3025 CST, DE, 1:250), anti-phospho-Akt (Ser473) (# 4060 CST, DE, 1:250), anti-total Akt (#4685, CS, DE 1:250). Secondary antibodies were anti-rabbit IgG HRP (CST, DE, #7074). For immunocytochemistry, quadriceps muscles were harvested and fixed in PFA (4%) for 72h and cryoprotected by further immersing in 30% sucrose solution for another 72 hr. After embedding in OCT, sections were cut and stained using standard immunohistochemistry using the following antibodies: anti-Dystrophin (#ab15277, Abcam, UK1:150). For the analysis of myofiber types, quadriceps muscles were harvested and frozen by immersing the unfixed quadriceps muscle in OCT and slowly freezing the block in pre-cooled isopentane solution in liquid nitrogen. Sections were cut and stained using standard immunohistochemistry using the following antibodies: BA-D5, SC-71 and BF-F3 (DSHB, Iowa, US 1:150) and counterstained with DAPI.

For analysis of mean fiber area, quadriceps cross-sectional area and mean muscular fiber number, dystrophin-stained muscle sections were drawn, segmented, and analyzed using Fiji software.

### Electron microcopy

Six month old animals were sacrificed, muscles were surgically excised, trimmed to small pieces and immediately immersed in cold 2.5% glutaraldehyde. Muscles were processed for electron microscopy as described previously (Takeshima et al., 2000). Briefly, tissue was cut into small pieces and fixed in 2.5% glutaraldehyde in 0.1 mol/l sodium phosphate buffer, pH 7.4. for 1 hour. Subsequently samples were rinsed with the same buffer and post-fixed in 2% osmium tetroxide in 0.1 mol/l sodium phosphate buffer on ice for 40 min. After 3 rinsing steps in ddH2O, the samples were dehydrated in a graded series of acetone on ice and embedded in Agar 100 resin. 70-nm sections were post-stained with 2% uranyl acetate and Reynolds lead citrate. Sections were examined with a FEI Morgagni 268D (FEI, Eindhoven, The Netherlands) operated at 80 kV. Images were acquired using an 11 megapixel Morada CCD camera (Olympus-SIS).

### Lipidomics

Quadriceps muscles were isolated from 8-week-old mice and snap frozen in liquid nitrogen. Muscle tissue was homogenized using a Precellys 24 tissue homogenizer (Precellys CK14 lysing kit, Bertin). Per mg tissue, 3μL of methanol were added. 20 μL of the homogenized tissue sample was transferred into a glass vial, into which 10 μL internal standard solution (SPLASH® Lipidomix®, Avanti Polar Lipids) and 120 μL methanol were added. After vortexing, 500 μL Methyl-tert-butyl ether (MTBE) were added and incubated in a shaker for 10 min at room temperature. Phase separation was performed by adding 145 μL MS-grade water. After 10 min of incubation at room temperature, samples were centrifuged at 1000xg for 10min. An aliquot of 450 μL of the upper phase (organic) was collected and dried in a vacuum concentrator. The samples were reconstituted in 200 μL methanol and used for LC-MS analysis. The LC-MS analysis was performed using a Vanquish UHPLC system (Thermo Fisher Scientific) combined with an Orbitrap Fusion™ Lumos™ Tribrid™ mass spectrometer (Thermo Fisher Scientific). Lipid separation was performed by reversed phase chromatography employing an Accucore C18, 2.6 μm, 150 x 2 mm (Thermo Fisher Scientific) analytical column at a column temperature of 35 °C. As mobile phase A we used an acetonitrile/water (50/50, v/v) solution containing 10 mM ammonium formate and 0.1 % formic acid. Mobile phase B consisted of acetonitrile/isopropanol/water (10/88/2, v/v/v) containing 10 mM ammonium formate and 0.1% formic acid. The flow rate was set to 400 μL/min. A gradient of mobile phase B was applied to ensure optimal separation of the analyzed lipid species. The mass spectrometer was operated in ESI-positive and -negative mode, capillary voltage 3500 V (positive) and 3000 V (negative), vaporize temperature 320°C, ion transfer tube temperature 285oC, sheath gas 60 arbitrary units, aux gas 20 arbitrary units and sweep gas 1 arbitrary unit. The Orbitrap MS scan mode at 120000 mass resolution was employed for lipid detection. The scan range was set to 250-1200 m/z for both positive and negative ionization mode. The AGC target was set to 2.0e5 and the intensity threshold to 5.0e3. Data analysis was performed using the TraceFinder software (ThermoFisher Scientific). Lipidomics results from five biological replicates per group were analyzed. The amount of each lipid was calculated as concentration per mg of tissue for all lipid species, measured in a single biological replicate. Values were next averaged over the five biological replicates for control and MckCre-Gpcpd1 muscle samples, log2 transformed, and compared between the groups using lipidR software.

### Glycerophosphocholine and choline targeted metabolomics

Choline and choline glycerophosphate were quantified by reversed phase chromatography on-line coupled to mass spectrometry, injecting 1 μl of the methanolic serum extracts onto a Kinetex (Phenomenex) C18 column (100 Å, 150 x 2.1 mm) connected with the respective guard column. Metabolites were separated by employing a 7-minute-long linear gradient from 96% A (1 % acetonytrile, 0.1 % formic acid in water) to 80% B (0.1 % formic acid in acetonytrile) at a flow rate of 80 μl/min. On-line tandem mass spectrometry (LC-MS/MS) was performed using the selected reaction monitoring (SRM) mode of a TSQ Altis mass spectrometer (Thermo Fisher Scientific), with the transitions m/z 104.1 → m/z 60.1, CE=20 (choline) and m/z 258.1 → m/z 104.1, CE=20 in the positive ion mode. Additionally, several amino acids (serine, isoleucine, leucine, tryptophan and phenylalanine) were recorded by respective Selected Reaction Monitoring (SRM) as internal standards. Data interpretation was performed using TraceFinder (Thermo Fisher Scientific). Authentic metabolite standards were used for validating experimental retention times by standard addition.

### Osmolarity measurements

Quadriceps muscles were isolated from 8-week-old mice, snap frozen in liquid nitrogen, ground to a powder and weighed. Tissue lysates or cell lysates were prepared using metabolomics lysis buffer (20% water/40% Acetonytril/40% Methanol) to reduce the interference of proteins on the measurements. The final solution was mixed with deionized water (1:3 ratio) and measured using K-7400S Semi-Micro Osmometer (Knauer, DE). Obtained measurements were normalized to tissue weight.

### Statistical analysis

All data are expressed as mean +/- standard error of the mean (SEM). Statistical significance was tested by Student’s two tailed, unpaired t-test or two-way ANOVA followed by Sidak’s multiple comparison test. Correlation analysis was performed using Two-tailed Pearson correlation test. All figures and statistical analyses were generated using Prism 8 (GraphPad) or R. Details of the statistical tests used are stated in each figure legend. In all figures, statistical significance is represented as *P <0.05, **P <0.01, ***P <0.001, ****P <0.0001.

## RESULTS

### Aging affects the metabolic Gpcpd1-GPC pathway in the muscles

We first set out to investigate if Gpcpd1 mRNA expression and GPC metabolite levels change as a result of muscle ageing. We isolated skeletal muscles from adult (6 months) and old (24 months) mice. Gpcpd1 mRNA levels were markedly downregulated, while there was a highly significant accumulation of GPC in the aged muscles (Figure 1A, B). Further, we mined a recently reported multi-tissue age-dependent gene expression analysis in rats for potential changes in Gpcpd1 expression associated with ageing (Shavlakadze et al., 2019). Indeed, we found Gpcpd1 to be among the topmost significantly reduced genes in muscles with advanced age, displaying a high level of inverse correlation between Gpcpd1 mRNA levels and age (*r* = -0.8858; P<0.008) (Figure 1C). This reduction was only evident in the skeletal muscle, while other tissues tested did not show such age-related Gpcpd1 mRNA decline (Supplementary Figure 1). This data indicate that Gpcpd1 expression is downregulated in the aged muscle of rodents, with a significant increase in its catalytic substrate GPC.

**Figure 1.**
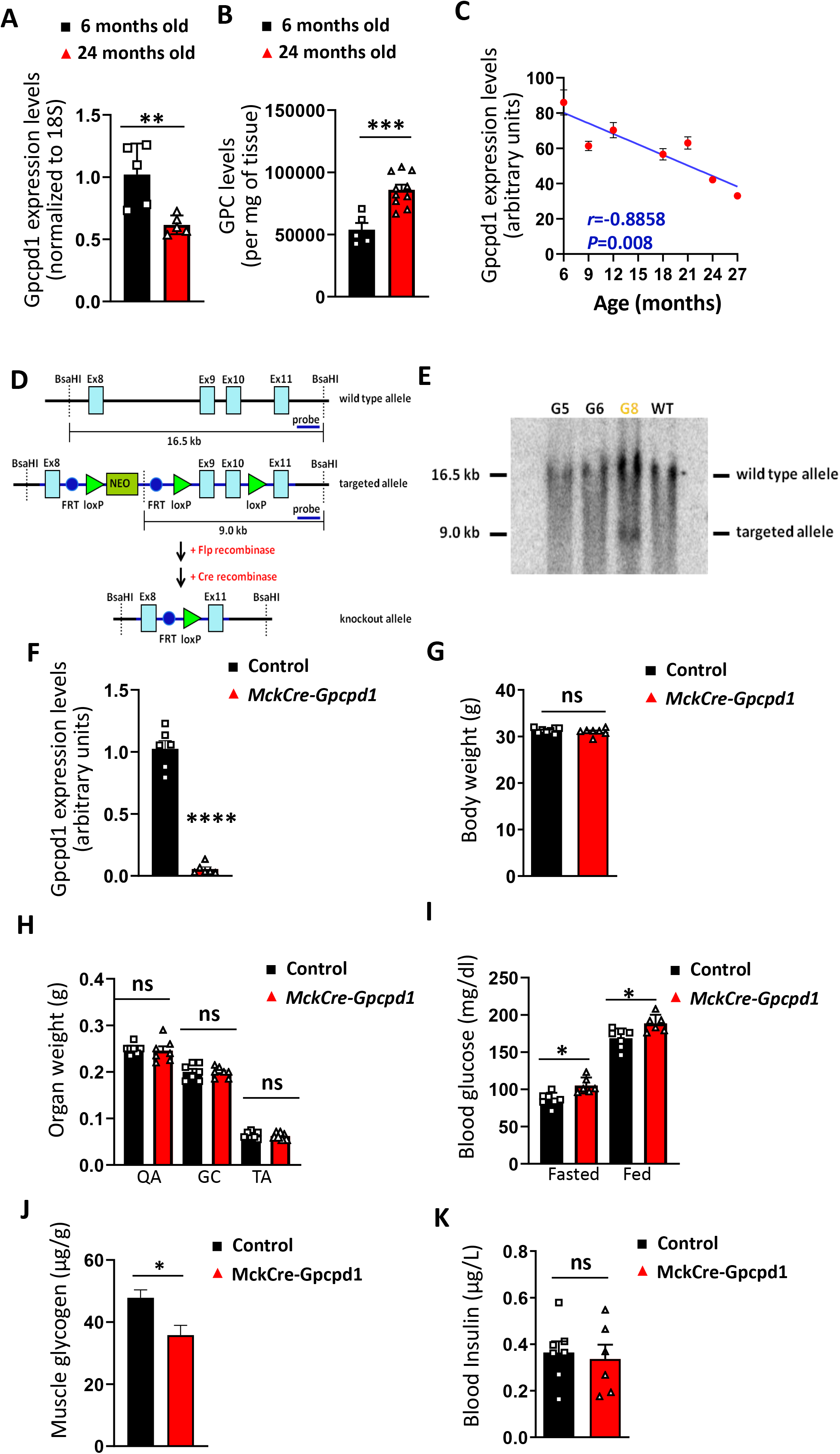
Muscle-specific *Gpcpd1* deficiency causes hyperglycemia in old mice. A) mRNA levels in quadriceps muscles of adult (6 months old) and old (24 months old) mice. N=5 per group. B) Glycerophosphocholine (GPC) levels in quadriceps muscles of young/adult (6 months old) and old (24 months old) mice. N=5-10 per group. C) Correlation of mRNA expression of Gpcpd1 in rat muscles with age. D) Schematic outline and E) Southern blot validation of successful generation of a conditional *Gpcpd1* allele in mice. F) Validation of Gpcpd1 mRNA deletion in muscles from muscle-specific *Gpcpd1* mutant mice (*MckCre-Gpcpd1^flox/flox^*). N=6 per group. G) Bodyweight of 20 months old control and littermate *MckCre-Gpcpd1^flox/flox^*mice. N=7-8 per group. H) Weights of skeletal muscle (quadriceps, QA; gastrocnemicus, GC; and Tibialis anterior, TA) isolated from 20 months old control and *Mck-CreGpcpd^flox/flox^* mice. N=7 per group. I) Blood glucose levels of 20 months old littermate control and *MckCre-Gpcpd1^flox/flox^*mice under fasted and re-fed states (2h after re-feeding; standard diet). N=6-7 per group. J) Skeletal muscle (quadriceps) glycogen levels in 20 months old control and *Mck-CreGpcpd1^flox/flox^* mice. N=6. K) Blood insulin levels of standard diet fed 20 months old control and *MckCre-Gpcpd1^flox/flox^*mice. N=6-7 per group. Unless otherwise stated, each dot represents an individual mouse. Data are shown as means ± SEM. Student’s two tailed, unpaired t-test was used for statistical analysis; ns, not significant; *p < 0.05; **p < 0.01; ***p < 0.001, ****p<0.0001.

### Muscle specific *Gpcpd1* deficiency does not affect muscle development but causes hyperglycemia in aged mice

To assess the physiological function of Gpcpd1 *in vivo,* we generated *Gpcpd1^flox/flox^* mice to specifically delete Gpcpd1 in defined metabolically important tissues that display a high Gpcpd1 expression (Figure 1D; Supplementary Figure 2). Successful gene targeting was confirmed by Southern Blot analysis (Figure 1E). As we found that Gpcpd1 expression was strongly reduced in aged muscles, we first crossed *Gpcpd1^flox/flox^* and *MckCre* transgenic mice to generate muscle-specific *Gpcpd1* deficient offspring, here termed MckCre *-Gpcpd1*^flox/flox^ mice. The efficiency of the *Gpcpd1* deletion in the muscle of *MckCre-Gpcpd1^flox/flox^* animals was evaluated by RT-PCR and by mass spectrometry (Figure 1F, Supplementary Figure 3A). Gpcpd1 mRNA levels were significantly reduced, with some residual expression most likely resulting from non-myogenic cells. As MckCre transgene is also active in the heart muscle(Bruning and Michael, 1998), Gpcpd1 was also deleted in the heart tissue of MckCre *- Gpcpd1*^flox/flox^ mice (Supplementary Figure 3B), while the expression in white adipose tissue (epididymal and subcutaneous fat), brown fat and liver was unaffected (Supplementary Figure 3C). Contrary to a previous report suggesting that over-expression of a truncated version of Gpcpd1 results in muscle atrophy (Okazaki et al., 2010), *MckCre-Gpcpd1^flox/flox^*mice exhibited apparently normal muscle development, muscle morphology, and muscle sizes at 6 month old and 20 months of age (Figure 1G,H, Supplementary Figure 4A). Moreover, the distribution of fiber type, myofiber size, muscle and mitochondrial ultrastructure, and finally muscle strength of 20 months old *MckCre-Gpcpd1^flox/flox^*mice appeared comparable to control mice (Supplementary Figure 4B-G). Intriguingly, aged (20 months old) developed fed and fasting hyperglycemia (Figure 1I). At the same time, there was a significant reduction in muscle glycogen levels (Figure 1J). Blood insulin levels remained unchanged (Figure 1K). These results indicate that muscle specific loss of *Gpcpd1* has no apparent effect on muscle development or muscle mass in aging but results in a muscle-mediated perturbation in glucose metabolism.

### Muscle-specific *Gpcpd1* mutant mice display glucose intolerance

To directly address muscle glycemic control, we performed a glucose tolerance test in young (12-week-old) littermate control and *MckCre-Gpcpd1^flox/flox^*mice. In contrast to older mice (Figure 1I) young *MckCre-Gpcpd1*^flox/flox^ mice did not yet display fasting hyperglycemia (Supplementary Figure 5A). However, 12-week-old mutant mice already displayed a severe glucose intolerance (Figure 2A,B) and overall increased insulin release (Figure 2C). Glucose intolerance was apparent also in the aged (20 months old) MckCre *-Gpcpd1*^flox/flox^ mice (Supplementary Figure 5B,C). To trace which organ is responsible for this defect, we bolus-fed (12-week-old) control and *MckCre-Gpcpd1*^flox/flox^ mice with radioactive 2-deoxy-D-glucose (2DG) and measured the uptake in several tissues. 2DG uptake was significantly reduced in the skeletal muscle (Figure 2D,E) and in the heart muscle (Supplementary Figure 5D,E) By contrast, 2DG uptake in white fat, subcutaneous fat, brown fat and liver was unaffected in *MckCre-Gpcpd1*^flox/flox^ mice (Supplementary Figure 5F). To test whether Gpcpd1 might also play a role in other metabolically relevant tissues that show high Gpcpd1 expression (Supplementary Figure 2), we crossed the *Gpcpd1*^flox/flox^ mice to mouse lines that mediate deletion in the liver, both white and brown fat, or the brown fat only. However, 12-week-old liver Gpcpd1-deficient (Figure 2F-H), white and brown fat Gpcpd1-deficient (Figure 2I-K), and brown fat Gpcpd1-deficient (Figure 2L-N) mice did not exhibit any apparent effects either on weight gain nor glucose metabolism, as addressed by glucose tolerance tests. These data show that muscle specific *Gpcpd1* deficiency results in glucose intolerance.

**Figure 2.**
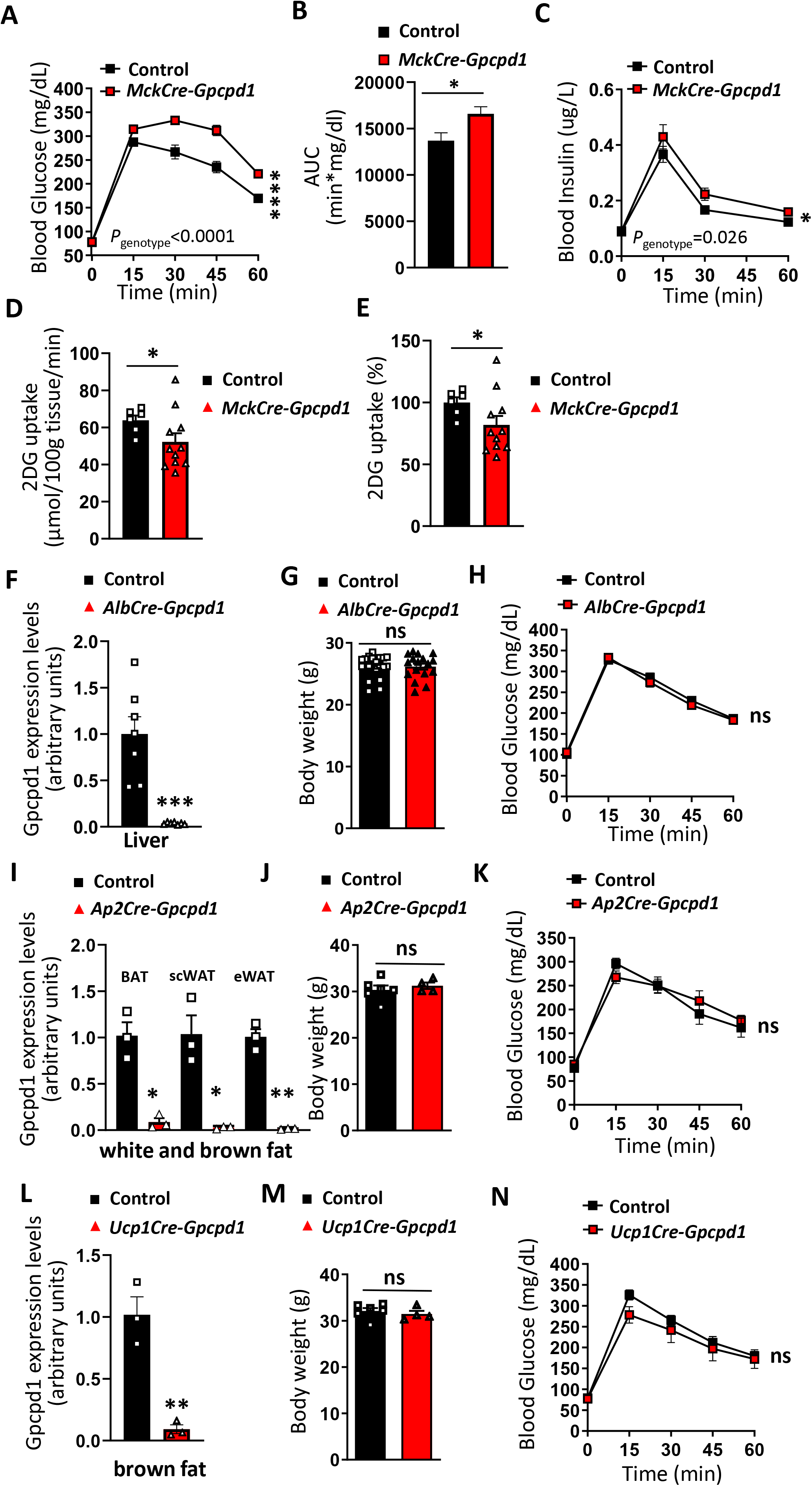
Muscle *Gpcpd1* deficiency causes glucose intolerance in young mice. A) Blood glucose levels and B) Area under curve (AUC) after an oral glucose tolerance test (OGTT) in 12 weeks old months old control and *MckCre-Gpcpd1^flox/flox^* mice. N=16-17 per group. Student’s two tailed un-paired t test with Welch correction was used for AUC statistical analysis. C) Blood insulin levels during OGTT in 12 weeks old control and *MckCre-Gpcpd1^flox/flox^* mice. N=12 per group. D) and E) Radioactive 2DG glucose uptake in skeletal muscles (soleus) of 12 weeks old control and *Mck-CreGpcpd1^flox/flox^* mice after bolus glucose feeding. F) RT-PCR confirmation of Gpcpd1 deletion in the liver of *AlbCre-Gpcpd1^flox/flox^* mice. G) Body weights of 3 months old control and hepatocyte-specific *Gpcpd1* deficient (*AlbCre-Gpcpd1^flox/flox^*) littermates. N=17-20 per group. H) Blood glucose levels after OGTT test in 2-3 months old control and *AlbCre-Gpcpd1^flox/flox^* mice. N=17-20 per group. I) RT-PCR confirmation of Gpcpd1 deletion in white fat (epididymal and subcutaneous) and brown fat of *Ap2Cre-Gpcpd1^flox/flox^* J) Body weights of 3 months old control and white and brown fat-specific *Gpcpd1* deficient (*Ap2Cre-Gpcpd1^flox/flox^*) littermates. N=4-6 per group. K) Blood glucose levels after OGTT in 2-3 months old control and *Ap2Cre-Gpcpd1^flox/flox^*mice. N=4-6 per group. L) RT-PCR confirmation of Gpcpd1 deletion in brown fat of *Ucp1Cre-Gpcpd1^flox/flox^* mice. M) Body weights of 3 months old control and brown fat-specific *Gpcpd1* deficient (*Ucp1Cre-Gpcpd1^flox/flox^*) littermates. N=4-6 per group. N) Blood glucose levels after OGTT test in 2-3 months old control and *Ucp1Cre-Gpcpd1^flox/flox^* mice. N=4-6 per group. Unless otherwise stated, each dot represents an individual mouse. Data are shown as means ± SEM. Repeated measures Two-way ANOVA followed by Sidak’s multiple comparison test was used for statistical analysis unless otherwise stated; ns, not significant; *p < 0.05; **p < 0.01; ****p < 0.0001.

### Muscle-specific *Gpcpd1* deficiency exacerbates high sugar and high fat diet induced metabolic syndrome

To further examine the metabolic dysfunction that develops in *MckCre-Gpcpd1^flox/flox^*mice, adult (3 months old) mice were first fed a high sugar diet (30% fructose in drinking water). There were no detectable differences in weight gain between controls and *MckCre-Gpcpd1^flox/flox^* mice over the 10 weeks observation period (Figure 3A). However, *MckCre-Gpcpd1^flox/flox^* mice displayed a severe defect in glucose clearance, as evaluated by a glucose tolerance test (Figure 3B,C). We next fed mice a high fat diet; again we observed no effects on weight gain but a severe glucose intolerance of the *MckCre-Gpcpd1^flox/flox^*mice as compared to their control littermates (Figure 3D,F). Insulin release appeared unaffected under both high sugar and high fat diet conditions (Figure 3G,H). Compared to the standard diet, high sugar and high fat diet tended to worsen the ability to regulate glucose metabolism of adult *MckCre-Gpcpd1^flox/flox^*mice (Figure 3I). These data show that inactivation of *Gpcpd1* in muscle does not affect weight gain under high sugar and fat diets but exacerbates glucose intolerance in adult *MckCre-Gpcpd1^flox/flox^* mice.

**Figure 3.**
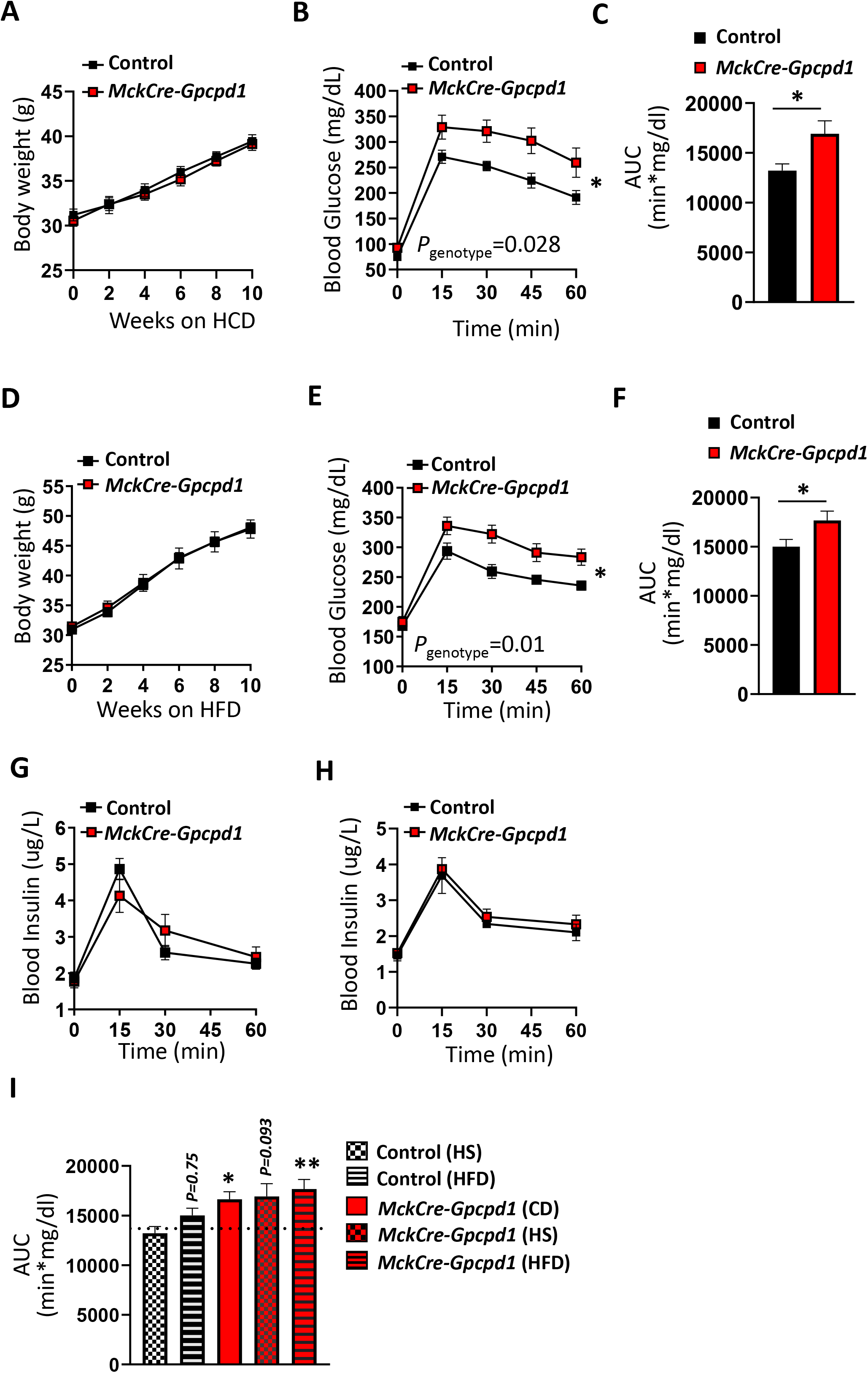
Muscle specific *Gpcpd1* deficiency exacerbates high sugar and high fat diet induced metabolic syndrome. A) Body weight gain of 3 months old control and *MckCre-Gpcpd1^flox/flox^*mice on high carbohydrate diet (HCD; 30% fructose). N=7 per group. B) Blood glucose levels after OGTT in and C) Area under curve (AUC) after 3 months old control and*MckCre-Gpcpd1^flox/flox^* mice fed a HCD diet. N=7 per group. Student’s two tailed un-paired t test with Welch correction was used for AUC statistical analysis. D) Body weight gain of 3 months old control and *MckCre-Gpcpd1^flox/flox^*mice on high fat diet (HFD; 60% fat). N=9 per group. E) Blood glucose levels after OGTT and F) Area under curve (AUC) after in 3 months old control and *MckCre-Gpcpd1^flox/flox^* littermate mice fed a HFD diet. N=9 per group. Student’s two tailed un-paired t test with Welch correction was used for AUC statistical analysis. F) Blood insulin levels during OGTT in 3 months old control and *MckCre-Gpcpd^flox/flox^* mice fed HCD. N=7 per group. G) Blood insulin levels during OGTT in 3 months old control and *MckCre-Gpcpd1* mice fed HFD. N=6 per group. H) Blood insulin levels during OGTT in 3 months old control and *MckCre-Gpcpd1* mice fed HFD. N=11-12 per group. I) Area under the curve (AUC) comparison during oral glucose tolerance test (OGTT) of 12 weeks old control and *Mck-CreGpcpd1^flox/flox^* mice that were fed either standard (chow diet), high sugar diet (HCD), and high fat diet (HFD). Multiple comparisons One-way ANOVA was used for statistical analysis. The effect of diets was compared to the AUC of control mice (dashed line) on standard diet after OGTT. Unless otherwise stated, each dot represents an individual mouse. Data are shown as means ± SEM. Repeated measures Two-way ANOVA followed by Sidak’s multiple comparison test was used for statistical analysis unless otherwise stated; ns, not significant; *p < 0.05, **p < 0.01.

### Muscle-specific *Gpcpd1* deficiency results in accumulation of glycerophosphocholine

Since Gpcpd1 is responsible for hydrolysis of glyceroposphocholine (GPC), we reasoned that muscle *Gpcpd1* deficiency would result in accumulation of GPC. To address this, we performed targeted lipidomic analysis on quadriceps muscle isolated from control and *MckCre-Gpcpd1^flox/flox^*mice. As expected, there was a significant accumulation of GPC in the muscle of *MckCre-Gpcpd1^flox/flox^* mice (Figure 4A). Besides the marked increase in GPC, there were no significant changes in the global lipidome in the muscles from *MckCre-Gpcpd1^flox/flox^* mice (Supplementary Figure 6A), indicating that loss of Gpcpd1 specifically impairs degradation of GPC. Choline levels remained unchanged (Supplementary Figure 6B), which could be explained because choline is mainly provided in the food (Fagone and Jackowski, 2013). Similar changes were observed in muscles isolated from aged (24 months old) mice when compared to adult (6 months old) wild type mice (Supplementary Figure 6C,D). To address whether solely the high concentrations of glycerophosphocholine can affect glucose uptake of muscle cells, we exposed C2C12-derived myotubes for several days with glycerophosphocholine (GPC) supplemented media. Using mass spectrometry, we confirmed that this treatment increased intracellular GPC levels in C2C12-derived myotubes (Figure 4H), with a trend towards increased intracellular osmolarity (Supplementary Figure 6E). Indeed, the treatment resulted in significantly reduced glucose uptake of C2C12-derived myotubes after 16h starvation (Supplementary Figure 6F). Thus increased concentration of osmolytes, such as GPC, inhibit glucose uptake in muscle cells. Taken together, inactivation of Gpcpd1 in the muscle results in marked and specific accumulation of the lipid GPC and a change in muscle osmolarity and glucose uptake.

**Figure 4.**
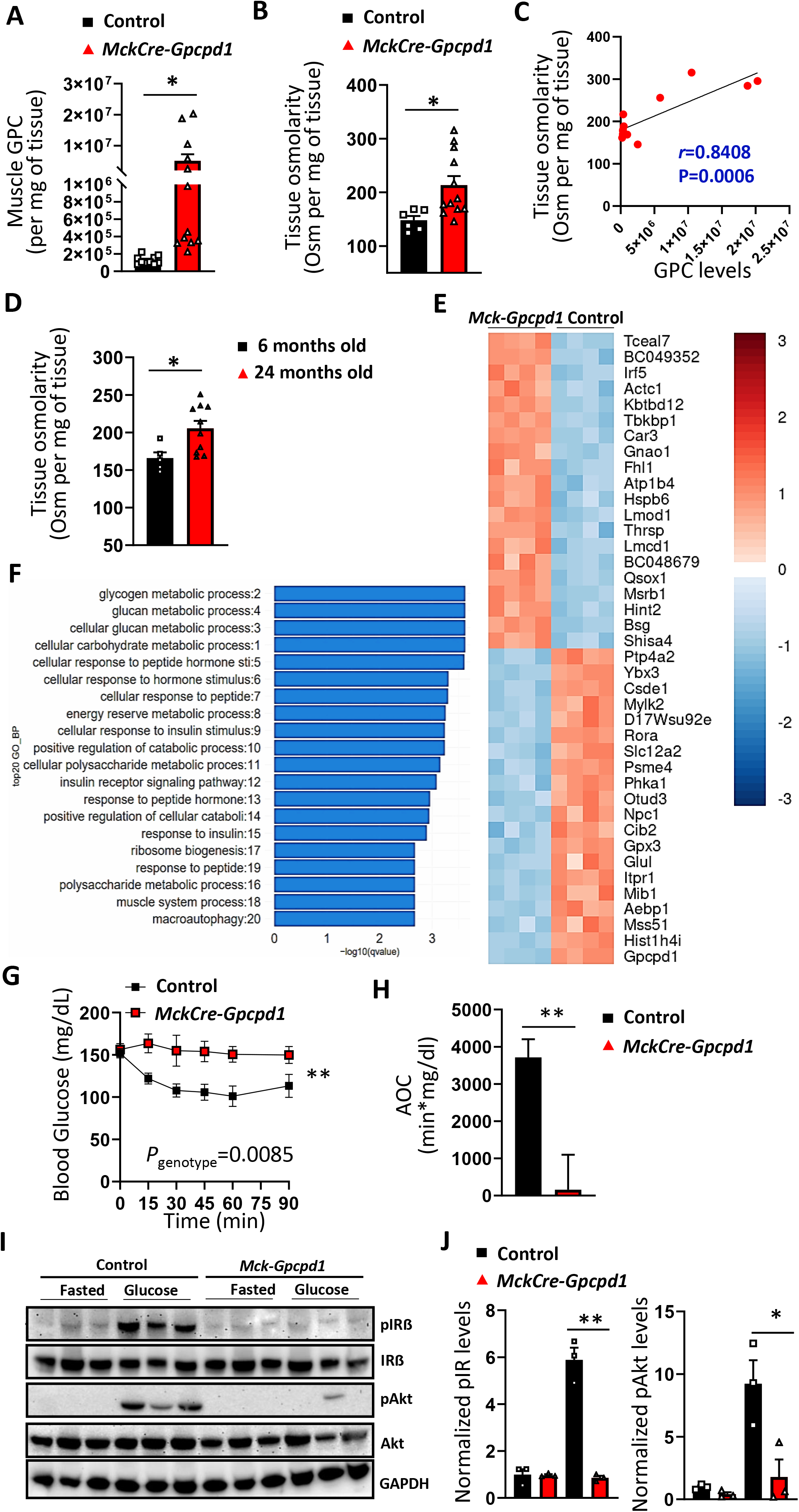
Muscle aging gene expression signatures and impaired Insulin receptor signaling upon Gpcpd1 loss in the skeletal muscle. A) Glycerophosphocholine (GPC) levels in skeletal muscle (quadriceps) of 3 months old control and *MckCre-Gpcpd1^flox/flox^*mice. N=10-12 per group. B) Skeletal muscle tissue osmolarity (quadriceps) of 3 months old control and *MckCre-Gpcpd1^flox/flox^* mice. N=6-7 per group. C) Correlation between tissue osmolarity and GPC levels in *MckCre-Gpcpd1^flox/flox^*mice. D) Skeletal muscle tissue osmolarity (quadriceps) of 6 months old and 24 months old wild type mice. N=5-10 per group. Each dot represents one individual animal. D) Quantseq transriptomic analysis in skeletal muscle (gastrocnemicus) of 3 months old control and *MckCre-Gpcpd1^flox/flox^*littermate mice. N=4 per group. E) Gene ontology analysis of down-regulated genes in skeletal muscle from 3 months old *MckCre-Gpcpd1^flox/flox^*mice, compared to control littermates. F) Insulin tolerance test and G) area of the curve (AOC) of 3 months old control and *MckCre-Gpcpd1^flox/flox^*mice. N=7-8 per group. Mice were fasted 6h before the test. Student’s two tailed un-paired t test with Welch correction was used for AUC statistical analysis. H) and I) Phsopho-Insulin receptor beta (IR β) and phospho-Akt levels in skeletal muscles (quadriceps) from 3 months old control and *MckCre-Gpcpd1^flox/flox^*mice after overnight fasting and 15 minutes after oral glucose delivery (bolus 1g glucose/kg of body weight). Total IR β levels, phosphorylated IR β (phosphoTyr1150/Tyr1151 IR β, total Akt and phosphorylated Akt are shown. GAPDH is shown as a loading control. N=3 mice per group, per treatment. Each dot represents one myofiber culture. Unless otherwise stated, each dot in A-C represents an individual mouse. Data are shown as means ± SEM. Repeated measures Two-way ANOVA followed by Sidak’s multiple comparison test and Student’s two tailed, Two-way Pearson correlation, unpaired t-test with Welch correction was used for statistical analysis; ns, not significant. *p < 0.05, ****p<0.0001.

### Muscle-specific *Gpcpd1* deficiency impairs insulin signaling

To obtain an overview of transcriptional changes occurring in muscles upon *Gpcpd1* deficiency, we performed genome wide mRNA-sequencing analysis (Quant-Seq) to compare the skeletal muscle gene expression profiles from young (12 weeks old) *MckCre-Gpcpd1* and control littermate mice (Figure 4E). Overall, 894 genes were significantly changed in skeletal muscles of *MckCre-Gpcpd1^flox/flox^*mice (401 down-regulated, 493 up-regulated; adjusted p-value < 0.01). Interestingly, a large set of significantly dysregulated genes (Figure 4E and Supplementary Figure 7) were also the top-most dysregulated genes previously associated with muscle ageing (Shavlakadze et al., 2019). This data indicates that, at least partially, young *MckCre-Gpcpd1^flox/flox^* mice display an” aged like” muscle transcriptional profile. Functional enrichment analysis of down-regulated genes in muscles of *MckCre-Gpcpd1^flox/flox^*mice revealed an enrichment for pathways linked to glucose and carbohydrate metabolism, and insulin signaling (adjusted p-value < 0.05) (Figure 4F, Supplementary Figure 8). Indeed, *MckCre-Gpcpd1^flox/flox^* mice displayed reduced insulin sensitivity as determined by an insulin tolerance test (ITT) (Figure 4G,H). Moreover, upon glucose challenge *in vivo*, we observed impaired insulin signaling as addressed by decreased phosphorylation of the insulin receptor beta (IRβ) and Akt (Figure 4I,J). Thus, inactivation of *Gpcpd1* in the muscle results in altered gene expression profiles associated with metabolic syndrome and aging and impaired Insulin receptor signaling.

### Changes of GPCPD1 and GPC levels in aging and type 2 diabetes in humans

We finally investigated how our findings relate to human physiology and aging. We found that in the skeletal muscles of otherwise healthy, middle aged/old (49-70 years of age) individuals, there was a significant reduction in Gpcpd1 mRNA expression in two independent cohorts (Figure 5A,B). In parallel, there was a highly significant increase of GPC and GPC/PDE levels in skeletal muscles of sedentary aged individuals (Figure 5C). A strong positive correlation between GPC or GPC-PDE levels was only evident with chronological age (*r*=0.7023, *P*<0.0001 and *r*=0.7220, *P*<0.0001, respectively) (Figure 5D,E), while there was no correlation between GPC or GPC/PDE levels with increased fat mass, muscle mass and body mass index (Supplementary Figure 9). Thus, the muscle GPC and GPC-PDE levels gradually increase with advancing age. As we observed that deficiency of *Gpcpd1* in mice results in impaired glucose metabolism, we also addressed if hyperglycemia in type 2 diabetic patients is associated with perturbed GPC metabolism. Indeed, there was a significant accumulation of GPC and GPC-PDE in skeletal muscles of diabetic individuals compared to age, gender and BMI matched controls (Figure 5F), while several other metabolites remained unchanged (Figure 5G). Taken together, our data show that the GPCPD1-GPC metabolic pathway is dysregulated with aging in humans, and that GPC accumulates in muscles of type 2 diabetes patients.

**Figure 5.**
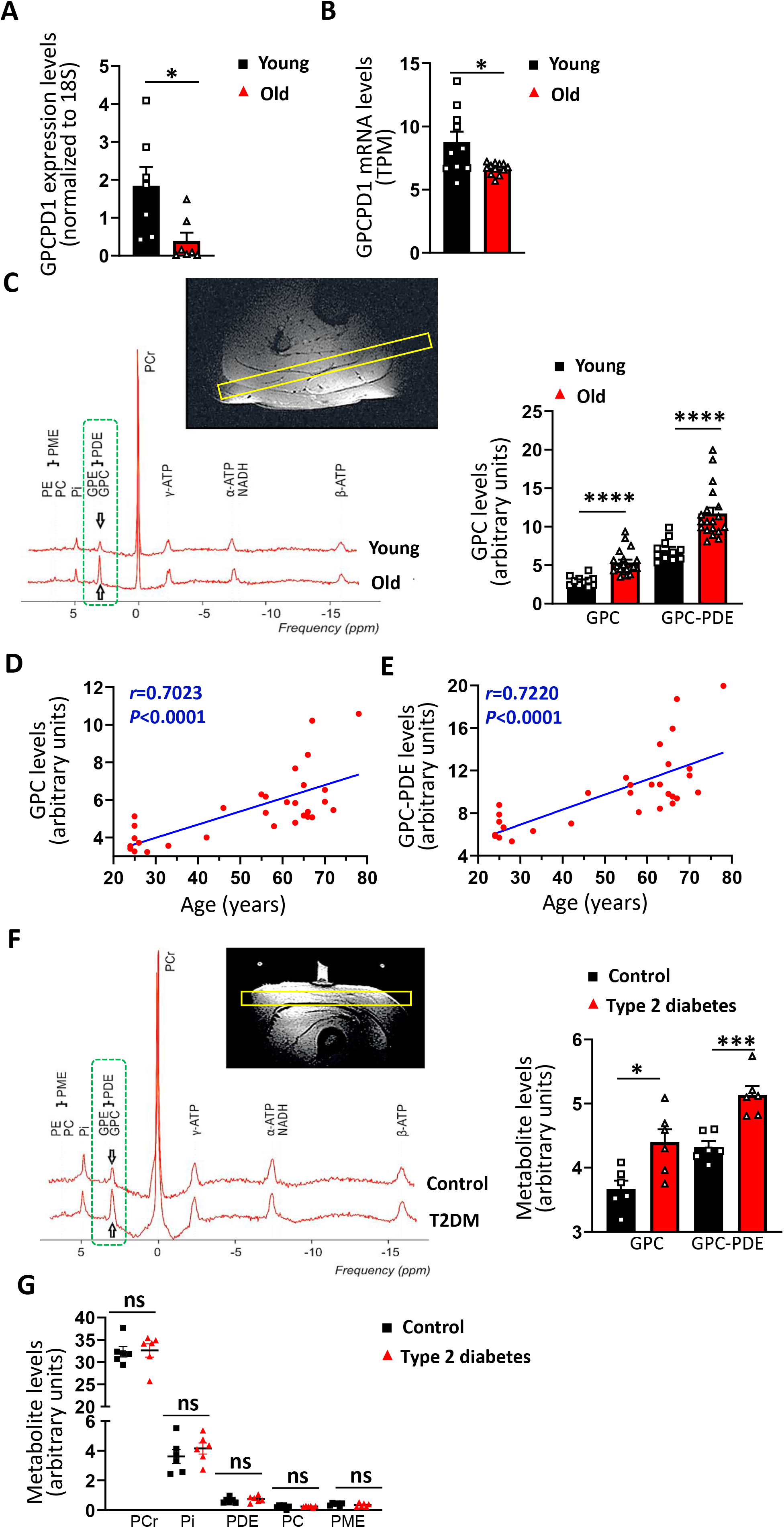
The metabolic GPCPD1-GPC axis is perturbed with aging and type 2 diabetes in humans. A) Gpcpd1 mRNA levels in quadriceps muscles in otherwise healthy young (20-30yrs) and old (49-65) humans and B) from Gpcpd1 quadriceps muscles in otherwise healthy young (25-35yrs) and old (60-70yrs) humans from two independent cohorts. Relative mRNA expression normalized to 18S RNA or transcripts per million (TPM) are shown. Relative mRNA expression normalized to 18S RNA is shown. B) Representative ^31^P-magnetic resonance (MR) spectra, and quantification of GPC and GPC-PDE levels acquired from the skeletal muscle (gastrocnemius medialis) from young (20-30yrs) and senior (49-65) participants. C) GPC and GPC-PDE levels plotted against age for each individual. Linear-regression analysis was used for R^2^ and P values calculation. E) Representative ^31^P-MR spectra, and quantification of GPC and GPC-PDE levels acquired from the skeletal muscle (gastrocnemius medialis) from healthy volunteers and type 2 diabetes patients (see Methods for cohort description). F) Levels of phosphocreatine (PCr), inorganic phosphate (Pi), phosphodiester (PDE), phosphocholine (PC), phosphomonoesthers (PME) acquired from the skeletal muscle from healthy volunteers and type 2 diabetes patients. Unless otherwise stated, each dot represents an individual human. Data are shown as means ± SEM. Student’s two tailed, Two-way Pearson correlation, unpaired t-test with Welch correction was used for statistical analysis unless otherwise stated; ns, not significant; *p < 0.05, ***p < 0.001, ****p < 0.0001.

## Discussion

Muscle aging is accompanied by a myriad of molecular and metabolic changes. Yet, it still remains elusive which of these perturbations are causative to the aging process, affecting both the muscle and systemic health (Demontis et al., 2013b). Here, we report that the Gpcpd1-GPC metabolic axis is perturbed with aging in muscles of rodents and humans. Based on these observations, we generated conditional and subsequently tissue specific *Gpcpd1* deficient mice to investigate if the observed age-related changes in this metabolic pathway affect systemic health. We found that muscle specific *Gpcpd1* deficiency results in a severe glucose intolerance, a disorder that is commonly seen in the elderly (Chia et al., 2018).

*Gpcpd1* deficiency has no apparent effect on body weight, muscle development, nor muscle mass even in old age. The apparently normal muscle mass and functionality in our model contrasts with a previous report where overexpression of a truncated Gpcpd1 protein *in vivo* resulted in muscle atrophy (Okazaki et al., 2010). This could be attributed to high levels of transgene expression, which can result in cell and tissue toxicity (Bolognesi and Lehner, 2018; Kulak et al., 2014; Moriya, 2015), not excluding other reasons for this previously observed phenotype such as dominant-negative effects of the overexpressed protein in defined pathways. We also deleted *Gpcpd1* specifically in several other metabolically important mouse tissues that exhibit Gpcpd1 expression, namely the liver and brown and white fat. However, we never observed any gross-developmental defects nor impairment in glucose metabolism. Thus, our results show that muscle-regulated glucose homeostasis is critically dependent on Gpcpd1 which could be, in part, explained by the relatively high Gpcpd1 expression in the muscles.

Both skeletal and heart muscle displayed impaired glucose uptake in the *MckCre-Gpcpd1^flox/flox^* mice. Given the relative organ to body weight, it is conceivable that the impaired glucose uptake in the skeletal muscle is mainly responsible for the systemic glucose metabolism perturbation in *MckCre-Gpcpd1^flox/flox^* mice. Future studies could reveal how Gpcpd1 loss only in the heart muscle affects heart function. Intriguingly, once we further generated Gpcpd1 deficient mice in several tissues with a relatively high Gpcpd1 expression and that are also essential to whole-body metabolic homeostasis, we did not observe any gross-developmental defects or impairment in glucose metabolism. Since Gpcpd1 has the highest expression levels in the striated muscles, we infer that the muscle tissue is specifically dependent on Gpcpd1 and its hydrolysis activity on GPC. A previous study has shown beneficial effects of GPC supplementation on the brain as well as cartilage tissue though the effect on the metabolic homeostasis was never tested (Matsubara et al., 2018). This could further indicate the muscle specific effects of the perturbation of the Gpcpd1-GPC metabolic pathway.

Mechanistically, our data show that *Gpcpd1* deficiency and increased GPC impaired muscle glucose uptake by interfering with glucose metabolism and insulin signaling. How would high levels of GPC, the specifically increased substrate of Gpcpd1, affect these pathways? It has been shown that the excess accumulation of other lipid species in the skeletal muscle, such as diacylglycerol, triacylglycerol, and ceramides, impairs insulin sensitivity and glucose uptake by perturbing mitochondrial beta oxidation and triggering an inflammatory program (Park and Seo, 2020; Wu and Ballantyne, 2017). It is possible that increased GPC level exerts similar effects, although we did not observe any signs of muscle inflammation in our *MckCre-Gpcpd1^flox/flox^*mice. At the transcriptional level, one of the most affected genes in the muscles from *MckCre-Gpcpd1^flox/flox^* mice in the insulin signaling network was Mss51 (Mitochondrial Translational Activator; also known as Zymnd17). Interestingly besides Gpcpd1, Mss51 is also one of the top-most downregulated genes in skeletal muscles with aging (Supplemental Figure 7) (Shavlakadze et al., 2019). Disruption of Mss51 in C2C12 using Cas9 system resulted in increased cellular ATP levels, upregulated glycolysis and oxidative phosphorylation (Rovira Gonzalez et al., 2019). However, *in vivo* disruption of Mss51 resulted in opposing effects in regard to glucose metabolism and insulin responses (Fujita et al., 2018; Rovira Gonzalez et al., 2019). *MckCre-Gpcpd1^flox/flox^*mice exhibit phenotype of severe glucose intolerance and impaired insulin signaling, without displaying any signs of mitochondrial dysfunction both at the transcriptional as well as ultrastructural levels. Further studies will be needed to elucidate which transcriptional or epigenetic mechanism is modulated due to Gpcpd1 loss and subsequent GPC accumulation. Interestingly, it has been previously reported that prolonged exposure to osmotic shock severely inhibits insulin stimulated glucose uptake and insulin signaling in adipose cells (Chen et al., 1999; Gual et al., 2003; Stookey et al., 2004). GPC acts as an osmolyte in kidney cells (Gallazzini et al., 2008). We detected increased osmolarity both in the muscle tissue of *Gpcpd1* mutant mice and in older wild type animals, while exposure of myofibers to GPC indeed impaired glucose transport. Therefore it is plausible that a chronic exposure to GPC changes this physicochemical property of muscle cells, contributing to the aging-induced impaired glucose homeostasis (Chia et al., 2018).

Importantly, muscles from aged humans and from patients with type 2 diabetes, displayed persistently increased levels of GPC, with a strong positive correlation between high muscle GPC levels and advancing age. Given our novel findings on the link between Gpcpd1-GPC metabolic pathway dysfunction and glucose intolerance, it is plausible that a chronic dysfunction of this pathway contributes to age-related impairment in glucose metabolism and diabetes (Chia et al., 2018). Diabetes is one of the leading chronic medical conditions in aged population, and a high risk factor for several age-related pathological conditions such as coronary artery disease, physical and cognitive decline, and mortality (Lee and Halter, 2017).

The large size of muscle tissue makes it an essential “metabolic sink” for glucose. Therefore, any age-related perturbation of this particular physiological function, strongly increases the risk of development of diabetes and consequently of all other age-related pathologies. Taken together, our findings highlight the key and previously unreported physiological role for the GPCPD1-GPC metabolic pathway. Aging results in the perturbation of this pathway specifically in the muscle tissue from rodents to humans. Genetical deletion of the Gpcpd1 enzyme only in the muscle tissues results in a severe glucose intolerance and impaired whole-body glucose metabolism, which was not observed for several organs that are of key importance to whole-body metabolic glucose control. Mechanistically, we find that specific accumulation of GPC due to loss of Gpcpd1 in the muscle perturbs insulin signaling, resulting with an “aged like” transcriptional profile. These data indicate that the perturbation of the Gpcpd1-GPC metabolic pathway may be involved in impaired glucose metabolism that is observed in the aged population. Moreover, there was a strong correlation of accumulating GPC levels with advanced age (r=0.72, *P*<0.0001), offering the possibility of using a non-invasive MRI-based readout of GPC as a muscle aging clock, without the need of muscle biopsies. Finally, whether this pathway can be efficiently therapeutically exploited to ameliorate the muscle-mediated dysregulation of metabolic glucose homeostasis in the elderly, needs to be further explored.

## Author contributions

D.C. and J.M.P. conceived, coordinated, and designed the study. D.C. and M.O. performed experiments and analyzed the data with contributions from M.L. and S.J.F. M.N. performed bioinformatic analysis. A.H. assisted in tissue sampling. E.R. and T.G. collected human muscle biopsies. L.P., R.K. and M.Krs. performed and analyzed in vivo MRS measurements. B.L., M.L., C.B., C.K., G.T., V.B., and C.M. performed glucose uptake experiments. N.V. generated and analyzed mRNA levels in the clinical cohort. G.D. and E.R. performed mass spectrometry experiments. M.L., M.K., M.Krs. and A.K.-W. designed, coordinated and oversaw the human T2DM experiments. D.C. and J.M.P. wrote the manuscript. All authors edited the manuscript and approved the final manuscript.

## Acknowledgements

We would like to thank all members of our laboratories for helpful discussions. We are grateful to Transgenic unit, Comparative medicine and Metabolomics unit from Vienna Biocenter Core Facilities for their service. We would also like to thank M.G. from the Histopathology unit in Vienna Biocenter Core Facilities for histological processing. We also than the lipidomic service of Center for Molecular Medicine (CEMM). We would like to thank Dr. Patrik Krumpolec from the Biomedical Research Center, Institute of Experimental Endocrinology, Slovak Academy of Sciences, Bratislava, Slovakia, for the processing of MRS data from young and senior participants. We would like to thank Prof. S. Trattnig, High Field MR Centre, Department of Biomedical Imaging and Image guided Therapy for logistic support. We would like to thank Prof. James D Johnson from University of British Columbia, Vancouver, Canada for critical reading of the manuscript.

J.M.P. is supported by IMBA, a Wittgenstein award, the T. von Zastrow foundation, and a Canada 150 Research Chair in functional genetics. D.C. is supported by the Austrian Academy of Sciences and the T. von Zastrow foundation. M. Krs., R. K. and L. P. are supported by Austrian Science Foundation (KLI-904-B).

## sSupplementary Figure legends

**Supplementary Figure 1.**
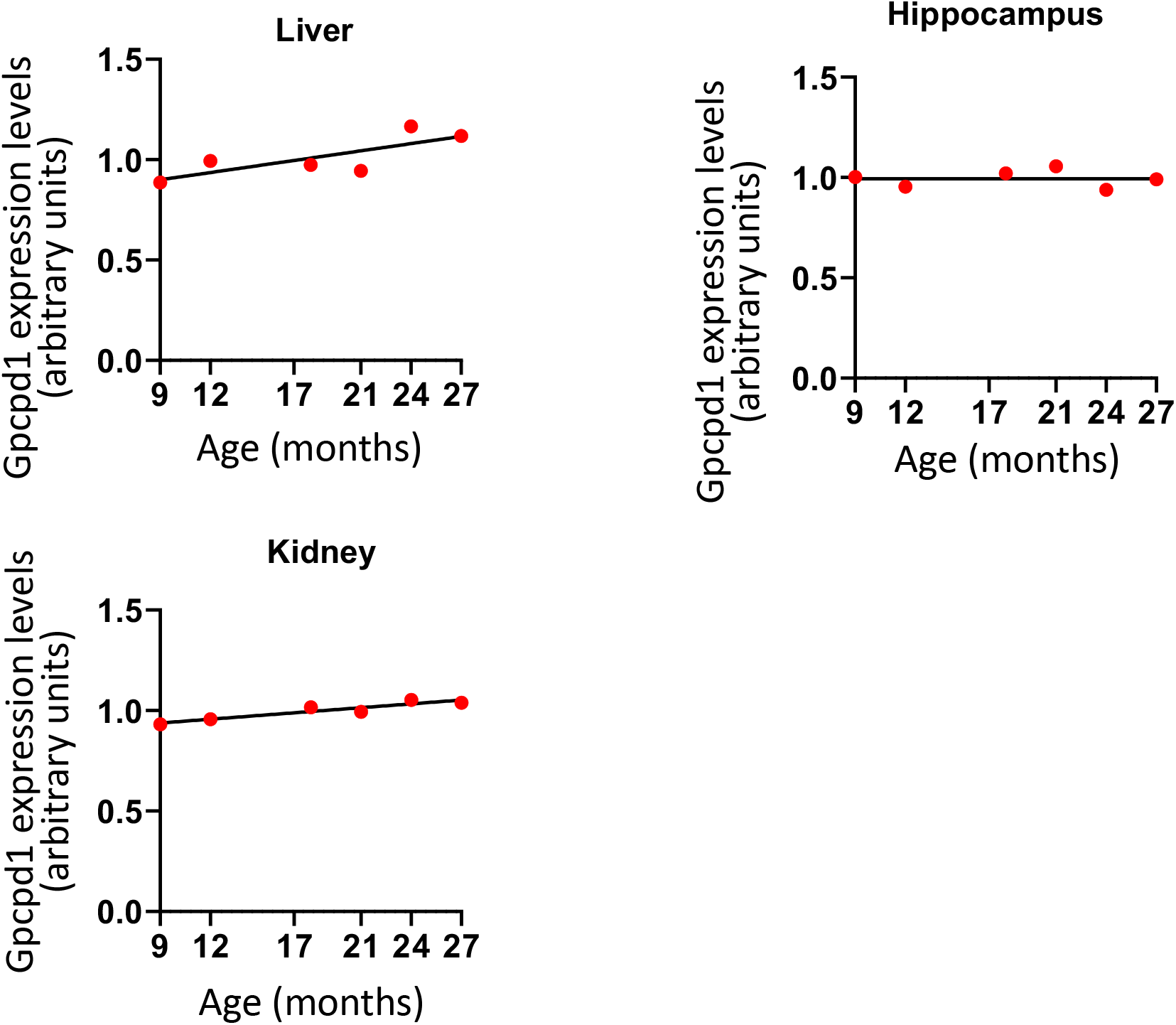
Assessment of age-related Gpcpd1 levels in rat tissues. Correlation of mRNA expression of Gpcpd1 in the indicated rat tissues with age. Data are shown as means ± SEM. Student’s two tailed, unpaired t-test was used for statistical analysis unless otherwise stated. There were no statistically significant differences.

**Supplementary Figure 2.**
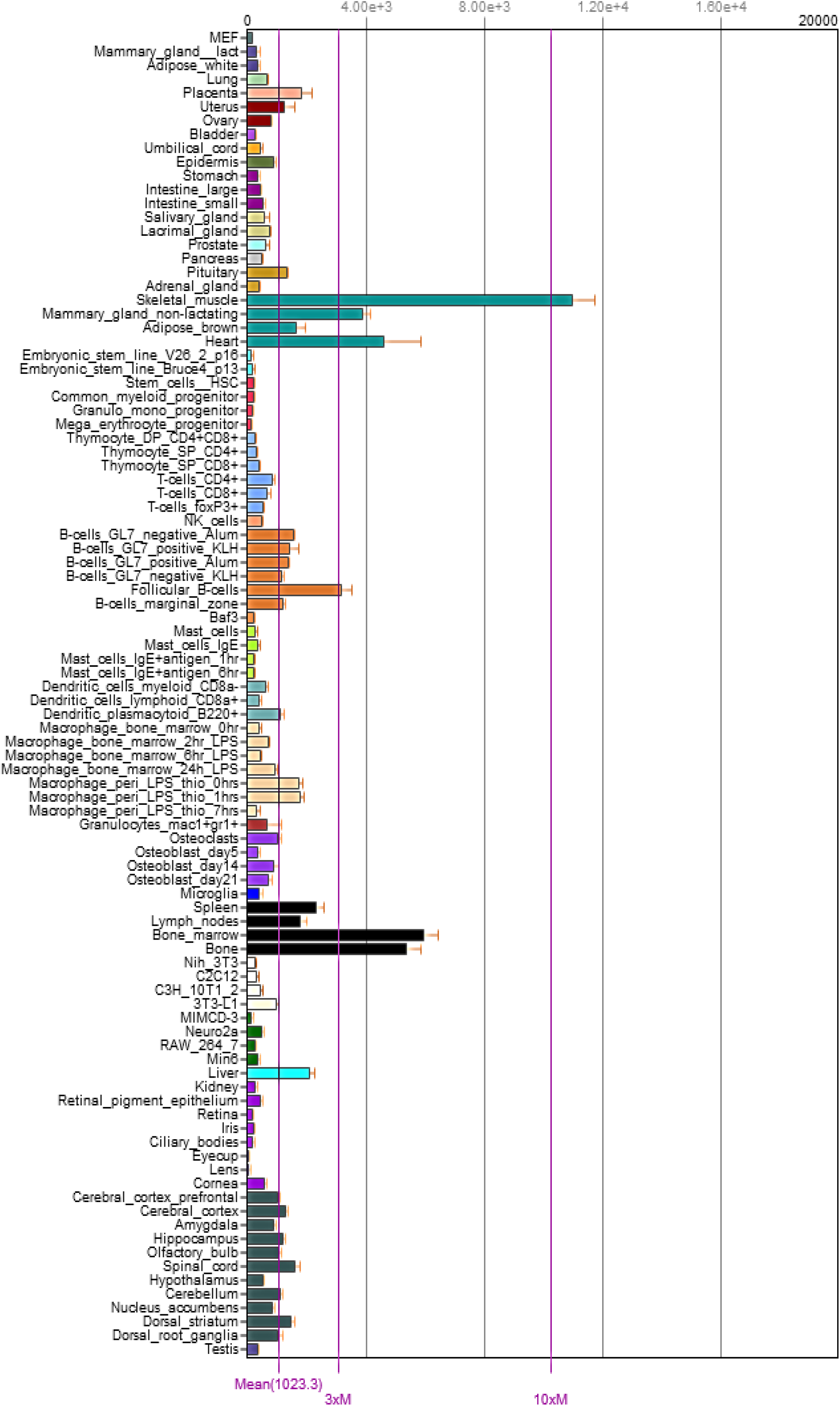
Gpcpd1 mRNA expression across different cell and tissue type. Gpcpd1 mRNA transcript per million in different tissues and cell types (http://biogps.org).

**Supplementary Figure 3.**
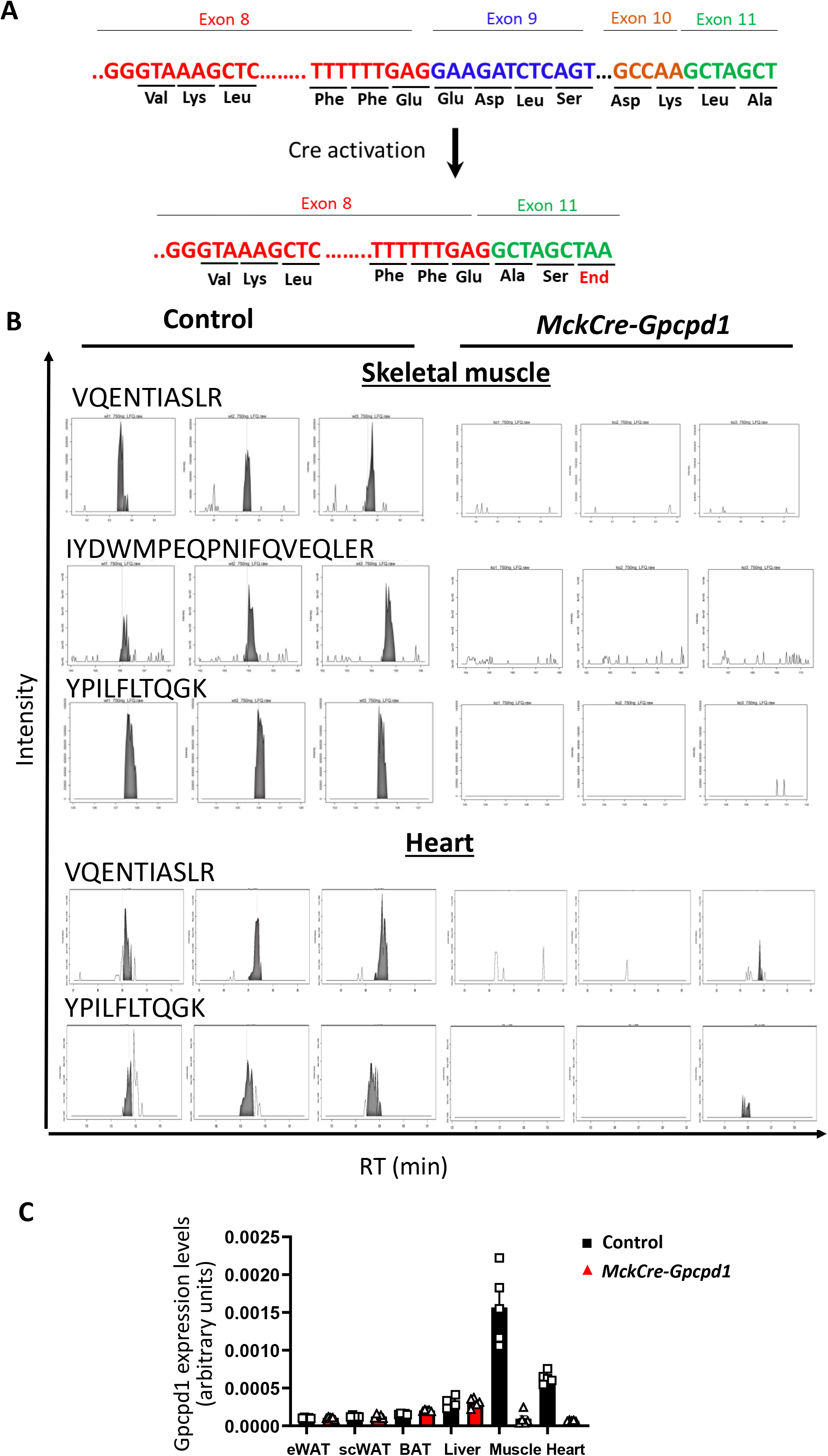
Validation of Gpcpd1 protein disruption in the skeletal muscle from *MckCre-Gpcpd1^flox/flox^*mice by mass spectrometry. A) Scheme of the KO strategy upon Cre activation B) Representative extracted ion chromatograms (XICs) of Gpcpd1 peptides downstream of exon 8 of the *Gpcpd1* locus from skeletal and heart muscle proteome of control and *MckCre-Gpcpd1^flox/flox^* mice. Integrated signal used for quantification is indicated in grey. Peptide identifications are represented by red dashed lines. XICs from N=3 mice per group are shown. C) RT-PCR based validation of Gpcpd1 deletion from various tissues in the *Mck-CreGpcpd1^flox/flox^* mice.

**Supplementary Figure 4.**
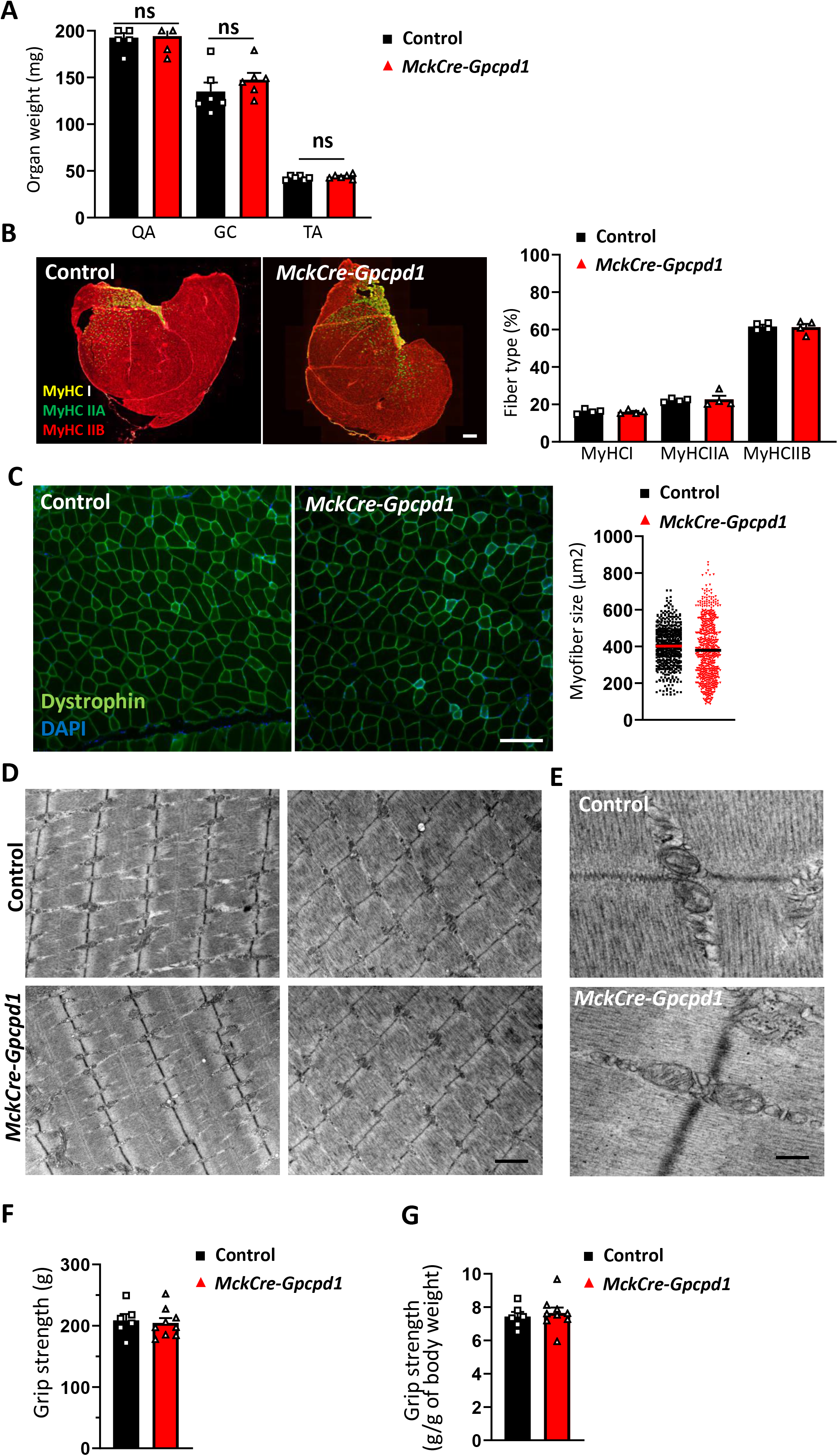
Grip strength, muscle weights, fiber type, and muscle ultrastructure assessment of *Mck-CreGpcpd1^flox/flox^*mice. A) Skeletal muscle weights (quadriceps, QA; gastrocnemicus, GC; and Tibialis anterior, TA) of 20 months old control and *Mck-CreGpcpd1^flox/flox^*mice N=6 per group, respectively. B) Representative images and quantification of MyHC!, MyhCIIA and MyHCIIB fibers in skeletal muscle (quadriceps) from 20 months old control and *Mck-CreGpcpd1^flox/flox^*mice. Images were taken under 5x magnification, and ≥100 myofibers were counted at 3 different matching histological areas. N=4 animals per group. Scale bar 500μm. C) Representative cross sections of skeletal muscle (quadriceps) and myofiber diameter size from 20 months old control and *Mck-CreGpcpd1^flox/flox^*mice. Myofibers were imaged using 10X magnification with ≥ 100 myofibers analyzed per mouse. n=4 animals per group. Scale bar 100μm. E) Ultrastructure of skeletal muscle (quadriceps) and F) intermyofibrillar mitochondria from 20 months old control and *Mck-CreGpcpd1^flox/flox^*mice. No apparent structural defects were observed in the muscles of *Mck-CreGpcpd1^flox/flox^*mice. n=3 animals per group. Scale bar=1μm and scale bar=500nm respectively. G,H) Muscle strength evaluation of 20 months old Control and MckCre-Gpcpd1 mice. Unless otherwise stated, each dot represents one individual animal. Data are shown as means ± SEM. Student’s two tailed, unpaired t-test with Welch correction was used for statistical analysis unless otherwise stated, ns, not significant.

**Supplementary Figure 5.**
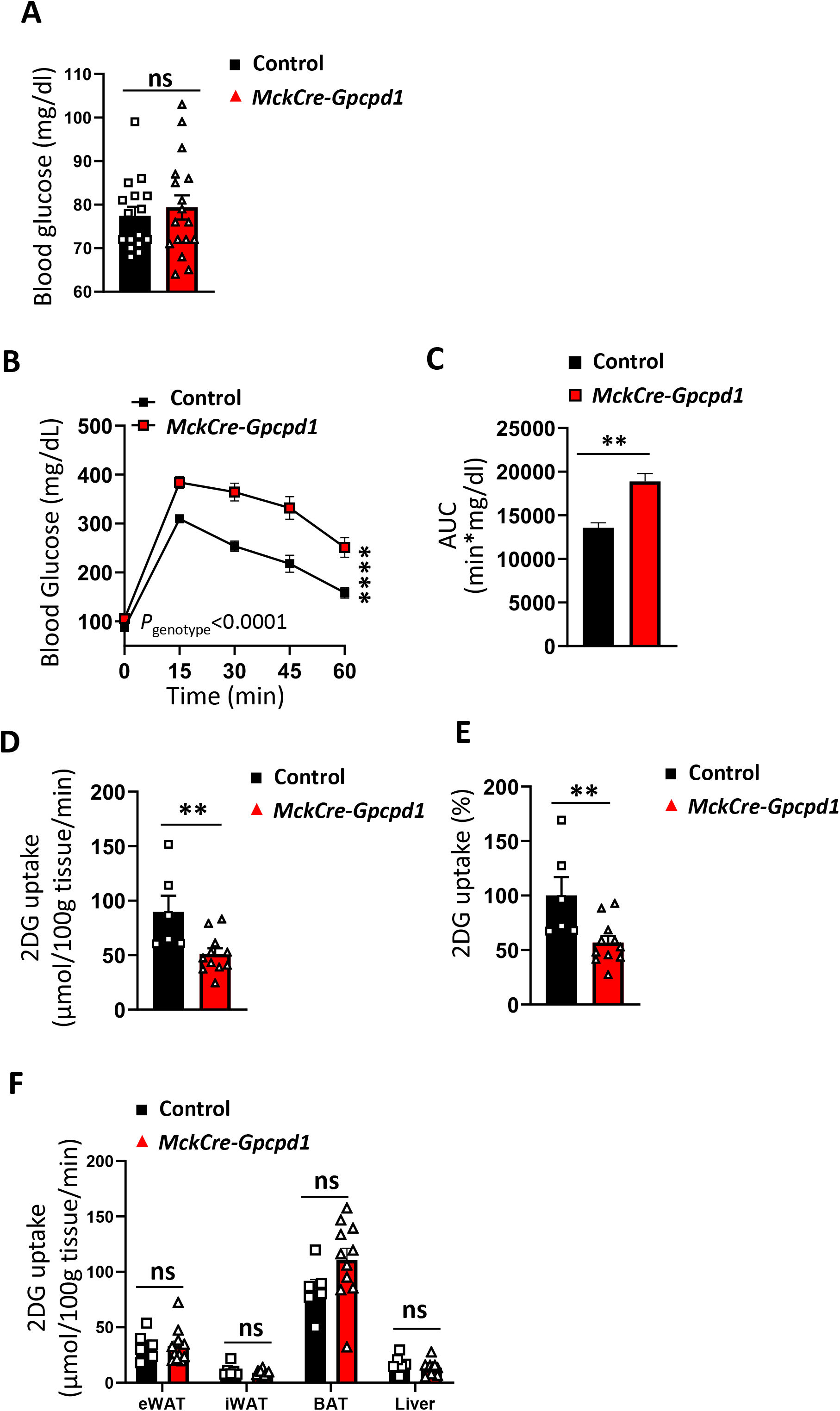
**Fasting glucose levels and tissue glucose uptake in *Mck-CreGpcpd1****^flox/flox^* **mice.** A) Fasting blood glucose levels in 12 weeks old control and *Mck-CreGpcpd1^flox/flox^*mice. N=6 and N=7 per group. B) Blood glucose levels and C) Area under curve (AUC) after an oral glucose tolerance test (OGTT) in 20 month old months old control and *MckCre-Gpcpd1^flox/flox^* mice. N=6 per group. Student’s two tailed un-paired t test with Welch correction was used for AUC statistical analysis. D) and E) Radioactive 2DG glucose uptake in heart muscle of 12 weeks old control and *Mck-CreGpcpd1^flox/flox^*mice after bolus glucose feeding. F) Radioactive 2DG glucose uptake in white fat (eWAT), beige fat (iWAT), brown fat (BAT), and liver of 12 weeks old control and *Mck-CreGpcpd1^flox/flox^*mice after bolus glucose feeding. Each dot represents one individual animal. Data are shown as means ± SEM. Student’s two tailed, unpaired t-test with Welch correction was used for statistical analysis unless otherwise stated; ns, not significant;*p < 0.05, ***p < 0.001.

**Supplementary Figure 6.**
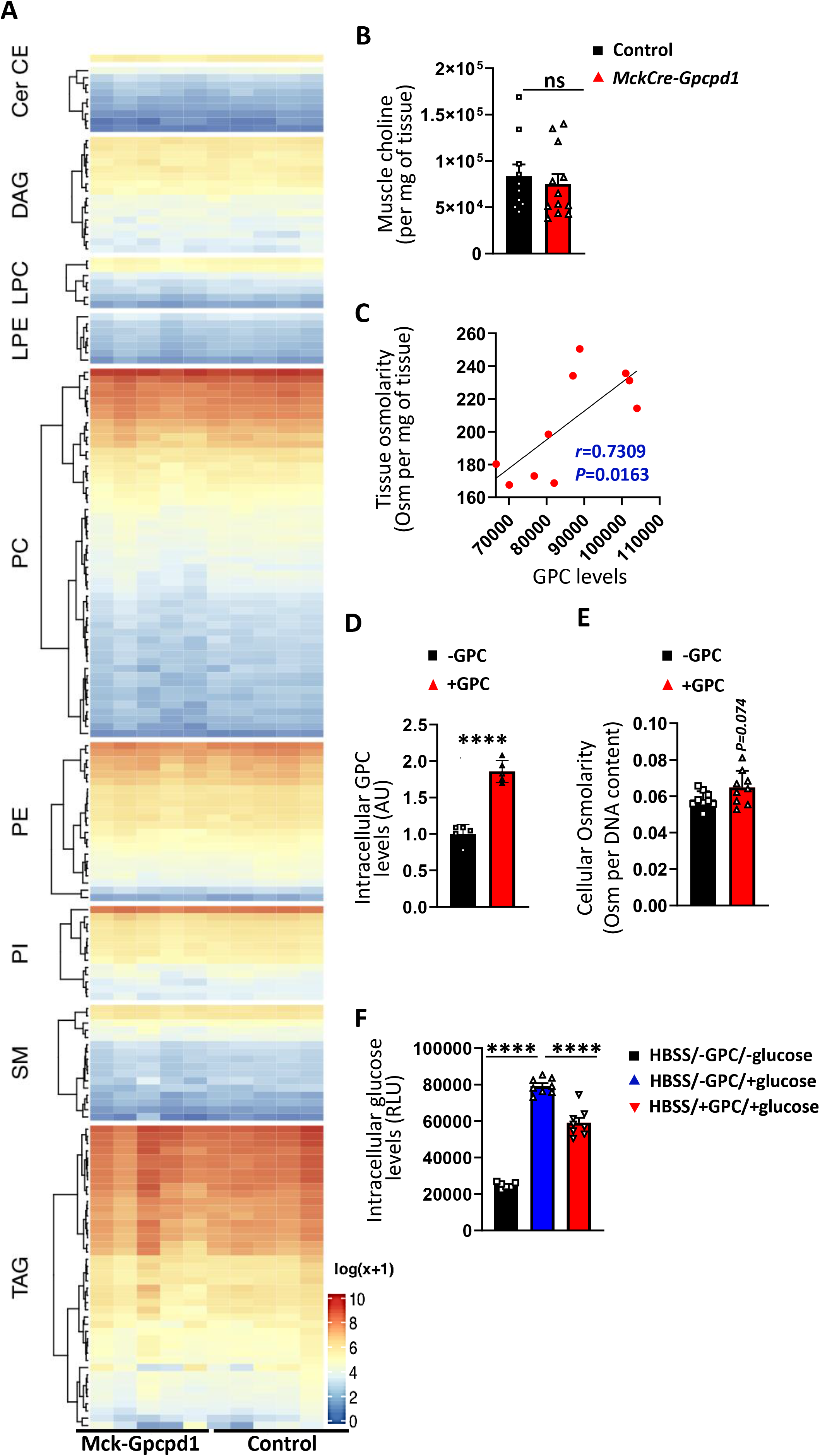
Muscle lipidome evaluation upon Gpcpd1 loss and C2C12 derived myotubes GPC treatment. A) Mass spectrometry based un-targeted lipidomic analysis in quadriceps muscles isolated from 3 months old control and *Mck-Cre-Gpcpd1 ^flox/flox^*littermate mice. Abbreviations denote triacylglycerols (TAG), sphingomyelins (SM), phosphatidylinositols (PI), phosphatydilethanolamines (PE), phosphatydilcholines (PC), lysophosphatydilethanolamines (LPE), lysophosphatydilcholines (LPC), diacylglycerols (DAG), ceramides (Cer), and cholesterolesther (CE). Numbers denote the carbon numbers, heatmap denotes higher and lower abundant lipid species in the muscles. No significant differences were found in the indicated lipid species abundance between muscles isolated from control and *Mck-Cre-Gpcpd1 ^flox/flox^* mice. N=5 per group. B) Choline levels in skeletal muscle (quadriceps) of 3 months old control and *MckCre-Gpcpd1^flox/flox^* mice. N=10-12 per group. C) Correlation between tissue osmolarity and rising GPC levels in young and old mice. Each dot represents individual mice. D) GPC levels in C2C12 differentiated myotubes without or with GPC pretreatment. Each dot represents one cell culture; error bars depict standard deviation. E) Cellular osmolarity of C2C12 derived myofibers treated with vehicle or GPC for 7 days ; error bars depict ; error bars depict standard deviation. F) Glucose uptake of C2C12 differentiated myotubes treated with vehicle or GPC for 7 days. Each dot represents one cell culture.; error bars depict Standard Deviation. Data are shown as means ± SEM unless stated otherwise. Student’s two tailed, unpaired t-test was used for statistical analysis unless otherwise stated, ns, not significant, ****p < 0.0001.

**Supplementary Figure 7.**
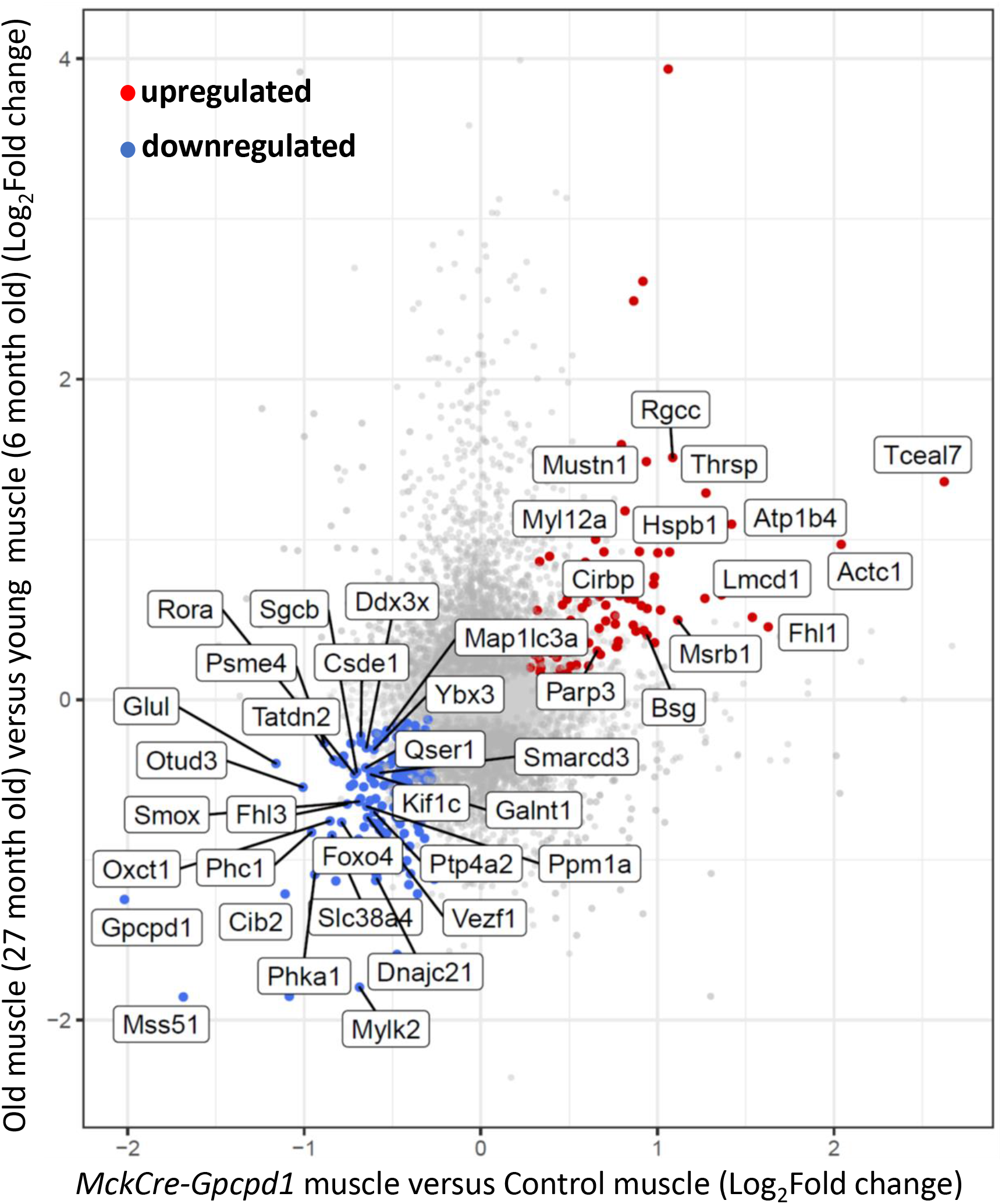
**Insulin signaling pathway s is affected in muscles from young *Mck-CreGpcpd1****^flox/flox^* **mice** Analysis of insulin signaling network on transcriptional level in quadriceps from young 3 months old control and *MckCre-Gpcpd1^flox/flox^* mice. The significantly downregulated genes (P<0,05) are labelled in green.

**Supplementary Figure 8.**
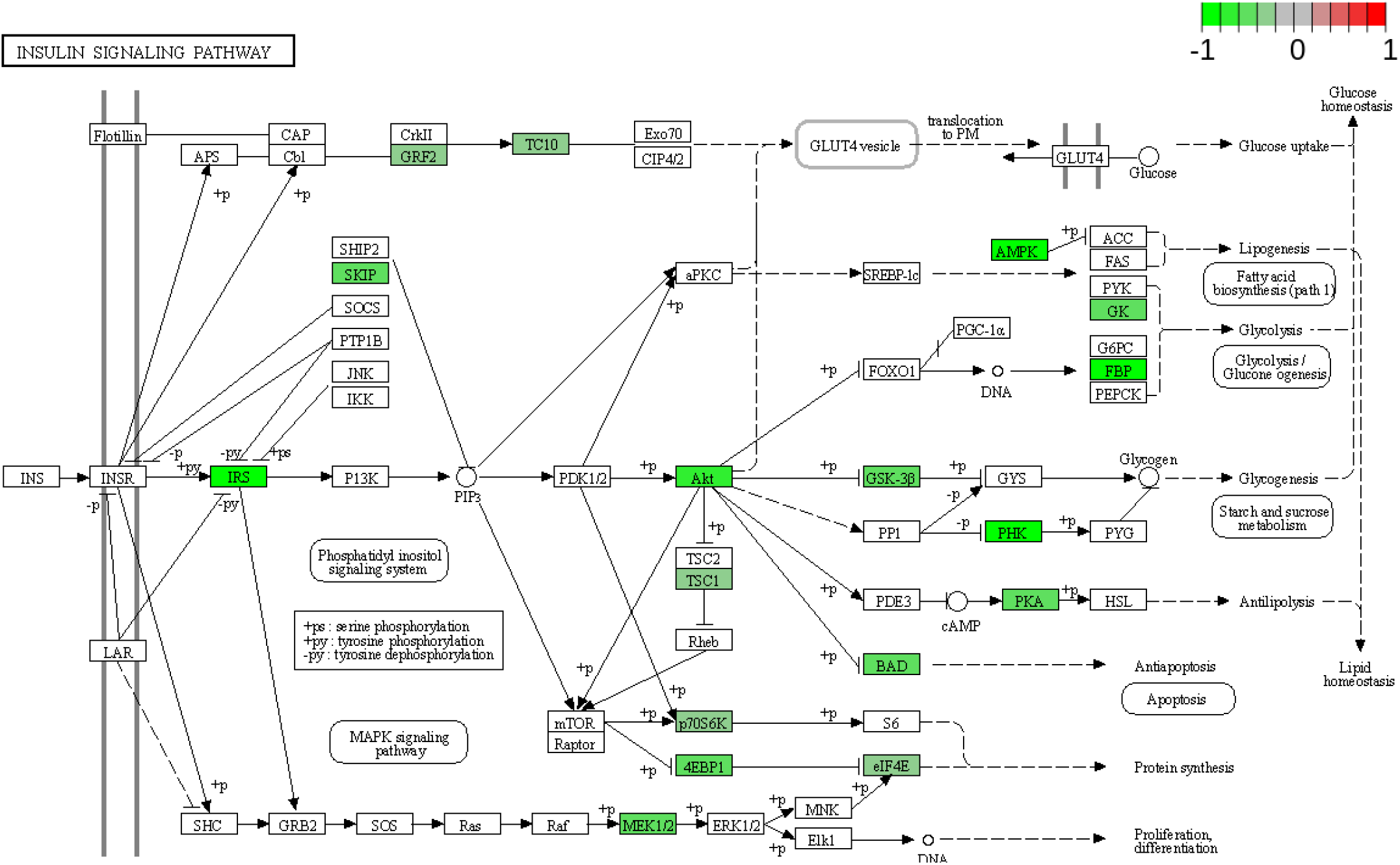
Muscles from young *Mck-CreGpcpd1^flox/flox^* mice display an “aged-like” transcriptomic signature. Comparison of upregulated (red) and downregulated (blue) gene expression changes in aged skeletal muscles in rats (27 month old versus 6 month old) (Shavlakadze et al., 2019) with gene expression changes of skeletal muscles from 3 month old control and Mck-CreGpcpd1^flox/flox^ mice. Overlapping dysregulated genes are highlighted.

**Supplementary Figure 9.**
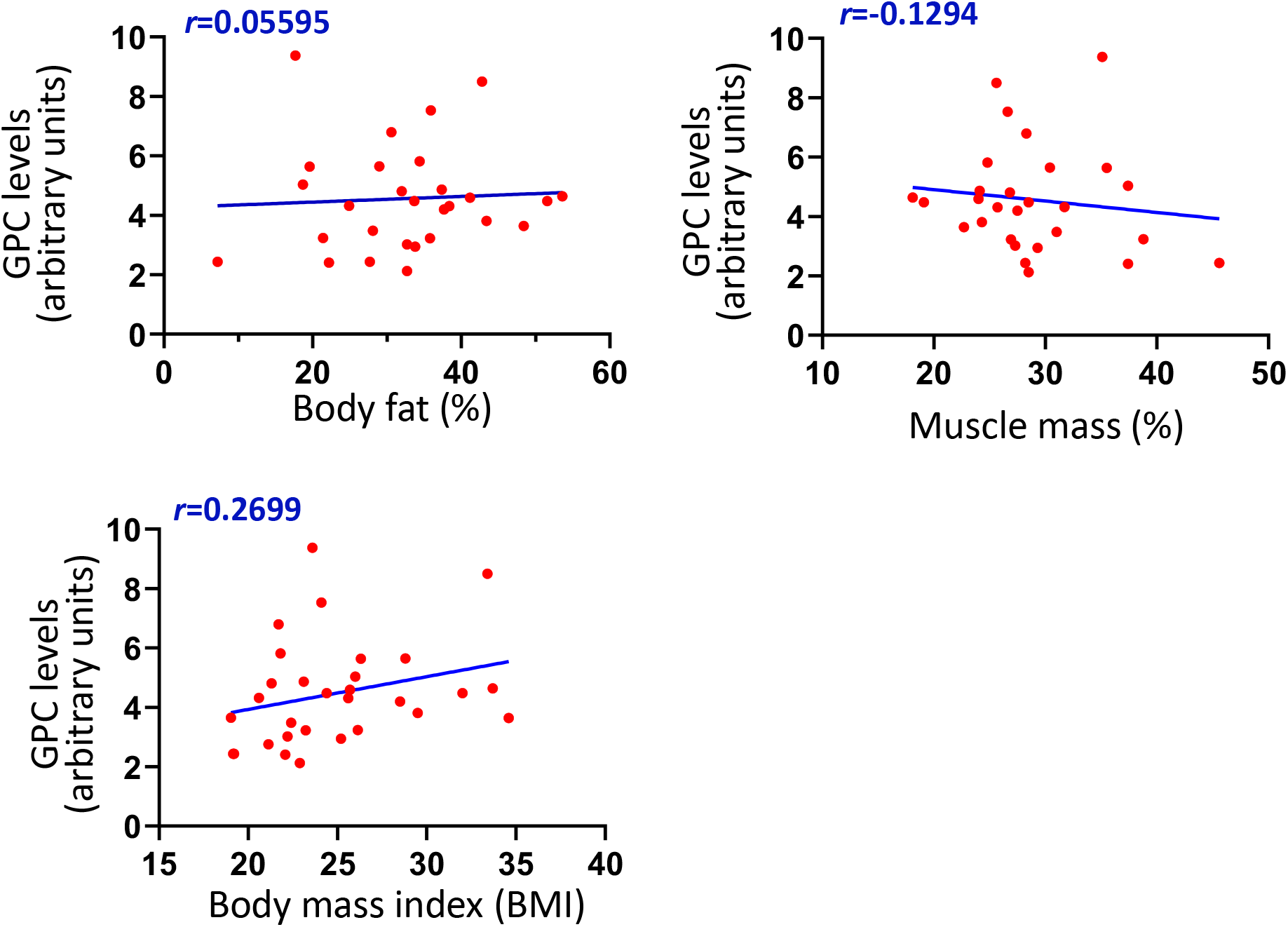
Correlation of GPC and GPC-PDE levels in humans with fat mass, muscle mass and body mass index. Correlation of skeletal muscle Glycerophosphocholine (GPC) and Glycerophosphocholine phosphodiester (GPC-PDE) levels with fat mass, muscle mass and body mass index in humans, irrespective of age. No significant correlation was found for any of the parameters. Two-tailed Person correlation test was used for the analysis.

